# Morpheus: A fragment-based algorithm to predict metamorphic behaviour in proteins across proteomes

**DOI:** 10.1101/2025.02.13.637956

**Authors:** Vijay Subramanian, Rajeswari Appadurai, Harikrishnan Venkatesh, Ashok Sekhar, Anand Srivastava

## Abstract

Functionally important “fold-switching” proteins, which do not obey the classical folding dogma, are now thought to be widespread. Algorithms that can accurately annotate fold-switching proteins from sequence information can help uncover the true extent of the “metamorphome”. Here, we present Morpheus, a fragment-based classification approach, that works by analysing the diversity of structures within a query protein sequence. Morpheus exhaustively curates and uses fragment structural data from the protein data bank as well as the AlphaFold Protein Structure Database. We employed our algorithm on 57 different proteomes consisting of a total of 601,218 proteins and identified about 10% of these proteins with the ability to fold switch. Additionally, we provide a web server for Morpheus to test for metamorphic propensities for user-defined sequences (http://mbu.iisc.ac.in/∼anand/morpheus). Besides screening for metamorphic behaviour in proteomes, our work will be useful in de novo design and engineering of such proteins through further experimentation.

## 1. Introduction

According to the ‘one-sequence, one-conformation’ paradigm proposed by Anfinsen [1], the amino acid sequence of a given protein codes for its three-dimensional folded structure required for the protein to perform its intended function. This paradigm has expanded to “one sequence, an ensemble of conformations” with highly labile but functional intrinsically disordered proteins (IDPs) [2]. Another branch of proteins that defy this paradigm, although not as conformationally degenerate as IDPs, is metamorphic proteins. Metamorphic proteins are a class of proteins that have the ability to exist in two conformations that are stable and significantly different in their secondary or tertiary structures [3, 4]. The two conformations correspond to highly dissimilar folds and possibly two different functions that the protein undertakes. Often, there is a biological trigger such as temperature, salt conditions, or the redox state associated with the conformational change that follows [5–8].

Some of the well-studied metamorphic proteins include the following: (a) RfaH, a virulence factor found in E. coli. RfaH is a multi-domain protein where the inter-domain interaction can modulate the conformation of the C terminal domain between an alpha-helical hairpin and a beta-barrel conformation. The all-alpha fold of RfaH functions to inhibit its activity, while the all-beta fold enables RfaH to activate transcription and facilitate translation [8–10]. (b) KaiB, the cyanobacterial protein switches fold to regulate the phosphorylation/ dephosphorylation phases of the circadian clock KaiABC, which allows it to maintain a periodicity of 24 hours [11]. (c) XCL1, the human lymphotactin is a metamorphic protein that undergoes a conformational rearrangement from a monomeric *α*+β chemokine fold to a novel dimeric all-β fold [12]. The canonical chemokine fold performs as an agonist for XCR1 (a G-protein coupled receptor), while the novel dimeric fold binds to surface glycosaminoglycans [13]. The trigger for the fold-switch in lymphotactin is temperature, and under physiological conditions, both folds are populated. Known metamorphic proteins are believed to be widespread in proteomes of many kingdoms of life and are currently estimated to constitute 0.5 - 4 % of the Protein Data Bank (PDB). [14]

Note the distinction between metamorphic proteins and fold-switching proteins that Porter and co-workers put forth: The main difference between metamorphic and fold-switching proteins is that the secondary structure remodelling events of fold-switchers can be either reversible or irreversible, but that of metamorphic proteins must always be reversible [15]. Hence, metamorphic proteins form a subset of fold-switching proteins. All the metamorphic proteins in the proteome can be called the “metamorphome.” Estimates have been made, but it is unclear how much of this class of proteins is present in the proteome overall. Protein sequences must be computationally screened for possible metamorphic behaviour and then validated using experimental methods in order to uncover the metamorphome in its entirety. Previous attempts have been made to predict metamorphic behaviour in proteins using protein sequence information. One of the approaches, from Chen and co-workers, is guided by the hypothesis that metamorphic proteins sequence would lead to degenerate secondary structure predictions [16]. Leveraging this degeneracy or confusion in predicting the secondary structure from the sequence data, the authors devised the diversity index, a metric to quantify the confusion of assigned secondary structures by structure prediction tools. Attempts have also been made to predict those fold-switching proteins that undergo alpha-helix to beta-sheet transition using sequence information alone to a high degree of accuracy [17]. This particular type of fold-switching protein, where there is a major switch from alpha-helical conformation to a beta-sheet, constitutes a relatively small portion of the known metamorphic proteins overall. Algorithms that tackled a parallel problem of predicting sequence-similar fold-switchers has also been developed [18]. Here, two different proteins with high levels of aligned similarity were assessed to see if they assumed the same fold or different ones. In contrast to fold-switching proteins, the sequence-similar fold-switching proteins described above remodel their secondary structures in response to mutations. Precise successive point mutations in protein sequences can lead to entirely different protein folds [19].

Another set of approaches that are designed to address a broader problem are methods that employ Alphafold suite of software to study conformational changes [20–24]. It is unclear whether the results obtained from Alphafold-based techniques are generative or based on memory; there is evidence supporting both sides of the argument [25–27]. When these methods are used in conjunction with high-throughput bioinformatic approaches, good results can be achieved compared to the results obtained individually. While it is worthwhile trying to predict metamorphic proteins using sequence information alone, the approach falls short at making proteome-level predictions due to a high false negativity rate. Methods involving biasing Alphafold to predict the alternate conformations of metamorphic proteins are often unreliable, resource intensive and of high time complexity. Here, we present Morpheus, a fast fragment-based predictor of metamorphic proteins that can achieve an average cross-validation accuracy of 84.7% with Matthew’s correlation coefficient of 0.70. To achieve this, we have implemented a Trie-based search to tremendously expedite the fragment picking performance for large proteomes. We employ our algorithm on 57 different proteomes and present the results here. Finally, we provide a web-server for using Morpheus to predict metamorphic protein sequences that were not covered in the dataset presented in this paper (web-server link: Morpheus). Additionally, from our database, we shortlist a few candidate metamorphic proteins that are involved in crucial biological processes and are highly likely to switch folds. These shortlisted proteins can be excellent candidates for further experimental work.

## 2. Methods

### 2.1 Training Dataset

The set of known fold-switching proteins and proteins that are highly likely to have a single fold is used as a training dataset. This data set has been borrowed from existing curated literature [14, 16]. The set is further filtered to remove structures flagged obsolete and structures that were later found not to be fold-switching due to erratum in structure. Additionally, the training dataset is expanded to include the designed metamorphic proteins as well [28]. In total, 189 fold-switching sequences and 198 monomorphic protein sequences are considered for the training dataset. Please see Tables S1 and S2 in the Supporting Information (SI). We chose to include fold-switching proteins for the construction of a metamorphic protein classifier for a couple of reasons: (a) There are only a handful of truly metamorphic proteins and it is unlikely to train a model on such limited data meaningfully. (b) Our model predicts those metamorphic proteins where there is a large change in the secondary structure of the two conformations with high confidence. Hence, when predictions are made, analysing the highly confident hits could help in searching for potential metamorphic proteins.

### 2.2 Fragment Picking

The long query sequence is first broken into fragments using a sliding window of sequence size 7. Each fragment from the query sequence is searched for identical fragments across a database of protein sequences containing around 200,000 redundant proteins from Protein Data Bank [29] and around 1.2 Million protein structure predictions from Alphafold2. For the proteins from PDB that contain multiple chains, each chain’s secondary structure is extracted and stored separately in the database. Given our fragment-picking database size, 7-mer fragments give the optimal number of hits while maintaining the secondary structure diversity of the hits. Once the hits are obtained, the secondary structure of the hits is extracted for the purpose of scoring. The hits are filtered to remove those fragments whose middle residue is an unobserved residue or has a local pLDDT score of less than 70 in the case of experimentally solved structures and computationally solved structures, respectively. The secondary structures of the proteins are assigned from the coordinates of the structures using DSSP [30]. As we perform the fragment search on a redundant PDB database, many of the hits occurring from different protein structures may all be pertaining to the same protein sequence. Hence, to eliminate the over-representation of proteins that have multiple redundant structures with the same sequence deposited in the PDB, we use a sequence clustering algorithm to cluster the database with a sequence identity threshold of 100%. Sequence clustering is performed using the fast and sensitive tool MMseqs2 [31]. All sequence alignments with sequence identity scores of 100% and covering at least 80% (-c 0.80) of both sequences are grouped in one cluster. When performing the fragment picking for a query sequence, multiple fragments may be obtained from different protein structures within the same cluster; to account for that, when secondary structure probabilities are calculated, each hit from a cluster is weighted by the inverse of the total number of hits from the same cluster.

The fragment search for individual proteins is performed by iterating through the lines of the database with a fast linear search in Java, matching fragment sequences using Regular expression (REGEX). When run on our system (16-core AMD Ryzen 7 5700, 64 GB RAM), the search for one fragment across the database is completed at an average of 700 milliseconds. For performing proteome-level searches, fragment picking is done with a Trie data structure implemented search using the Python module Pygtrie. Initially, the fragments that must be searched for in a proteome are stored in a Trie data structure. Once all the fragments are stored in the Trie, the database is iterated over once. When a fragment present in the Trie is found, the corresponding secondary structure information is stored as a leaf for that fragment in the Trie. Our system completes the Trie-implemented search for a proteome with 20,000 proteins within 2 hours.

Figure 1 shows the fragment picking schematic for one 7-mer fragment `"RLRSFTT"` from MAD2 protein. The different structures containing the fragment sequence are depicted in the circles, and one structure from each cluster is shown. By analysing the secondary structure of the middle residue in each of the hits obtained, the middle residue propensity is calculated and depicted as a pie graph.

**Figure 1.**
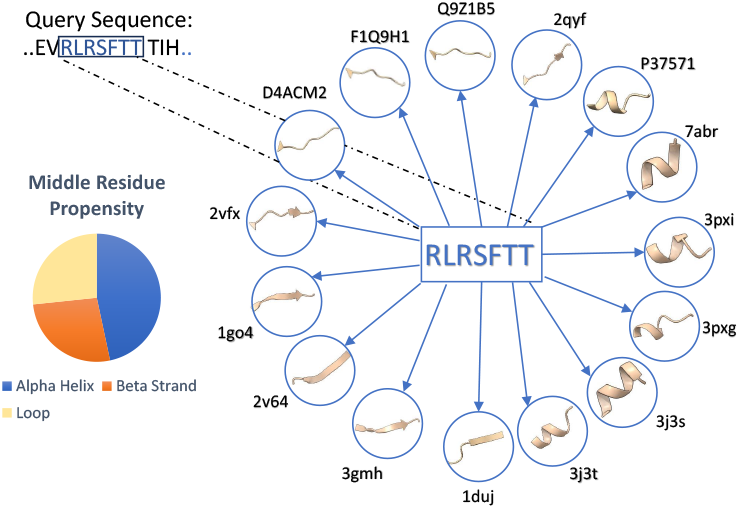
Fragment picking schematic. The query sequence fragment shown is a part of the metamorphic protein MAD2 and the cartoon representation of the 3D structures of proteins containing the fragments is shown in the circles. The PDB ID and the Alphafold Accession number are indicated in the representative hits. Figure S1 shows similar schematics for other fragment sequences taken from KaiB.

### 2.3 Scoring Metrics

A particular fragment is scored based on the different secondary structures of the hits obtained from the database. We hypothesized that local interactions within the fragment of 7 residues play a major role in determining the secondary structure of the middle residue. Fragment-based ab initio structure prediction tools like Rosetta [32] also choose to work with the secondary structure of the middle residue from the fragments that have been picked from their curated database for structure prediction. Guided by our hypothesis, we opted to analyse the secondary structure of the middle residue of the picked fragments. Therefore, the secondary structure of the middle residue is retrieved from each fragment hit obtained from our database. Since metamorphic proteins have the ability to inter-convert between two distinct conformations, often having a change in secondary structure, the metric we choose should be able to quantify the ability of a fragment sequence to adopt distinct secondary structures. Multiple consecutive fragments displaying this ability would, in turn, aid the whole protein sequence to switch folds. Following this thought, we consolidated four different metrics: **(a)** Diversity Index, **(b)** Substitution Score, **(c)** Information Entropy, and **(d)** Uncertainty Score. The diversity index [16] and substitution-matrix based scoring methods [14] have already been used for the classification of metamorphic proteins. Information entropy and uncertainty score are metrics that we have introduced for the purpose of classification of metamorphic proteins.

Diversity index and information entropy are defined as follows:

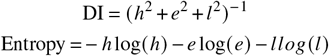

where h, e and l are the calculated propensities of the middle residue to be in helix, sheet and loop, respectively. The secondary structure propensities for each 7-mer are calculated from the hits obtained from the database by assessing what fraction of those hits have middle residues existing in helix, sheet and loop secondary structure. Hence, they are normalized and add up to 1. The diversity index and information entropy scores are calculated for all the fragments from their corresponding middle residue propensities. Figure 2 displays the diversity metrics for a few metamorphic and monomorphic proteins. Figure 2 shows that the region where the entropy score remains consistently high correlates well with the region of the protein that actually switches fold in the case of metamorphic proteins: KaiB, RfaH and Lymphotactin. As for the monomorphic proteins Ubiquitin, Galectin and Thermonuclease, the entropy scores remain low overall. Fig. S2 in SI portray the diversity metrics for other known metamorphic proteins such as RfaH, KaiB, IscU, MAD2, Lymphotactin, Selecase, MinE, CLIC1, HIV-RT1. We also show data for a designed protein Sa1V90T (PDB ID: 8E6Y) from John Orban and co-workers [28].

**Figure 2.**
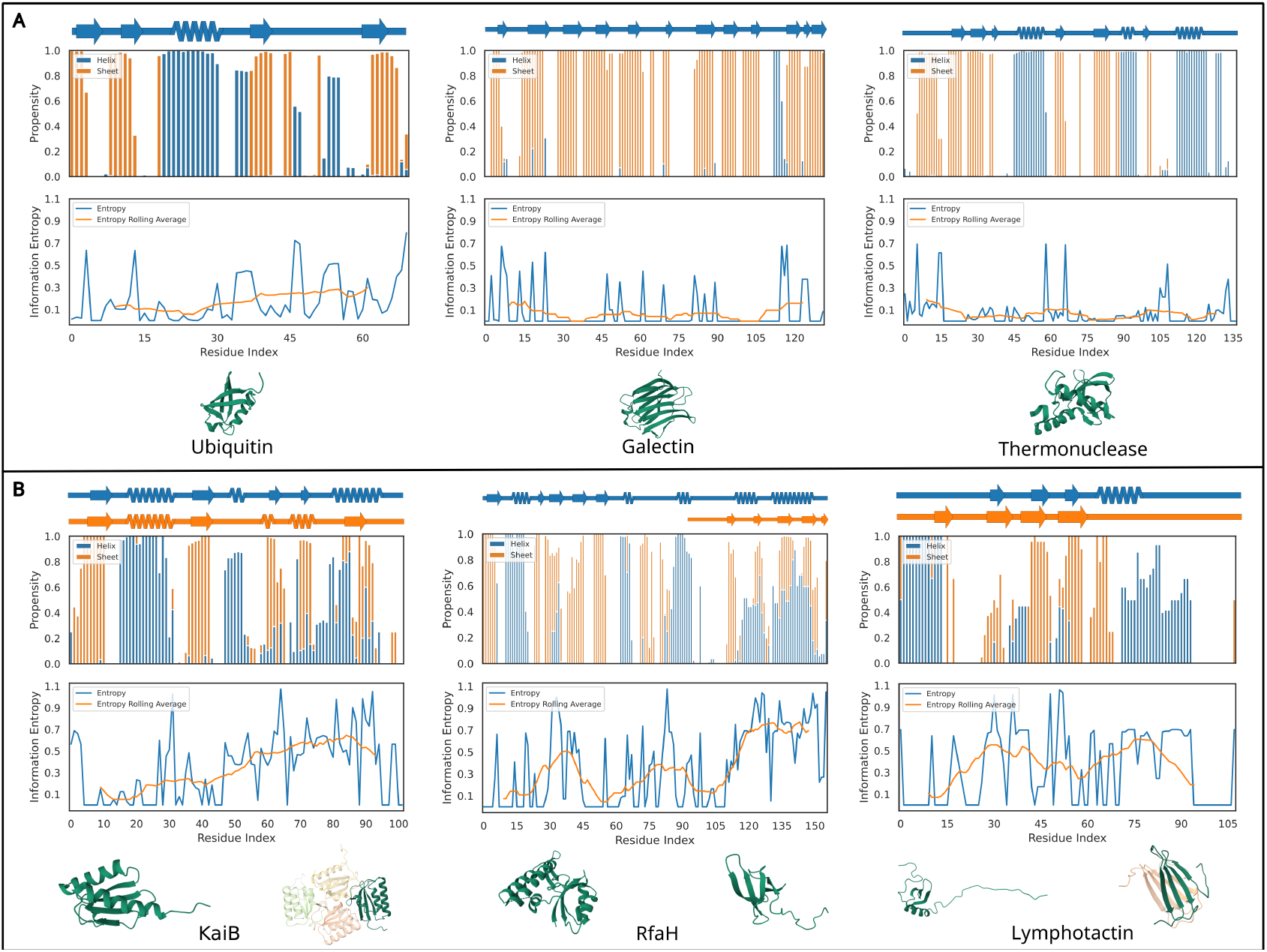
Diversity metrics able to distinguish metamorphic proteins and capture regions of fold-switching. The Secondary structure diagram annotated through an experimentally solved structure (top), secondary structure propensity plot (middle) calculated via fragment picking by Morpheus and entropy scores (bottom) are plotted for **(A)** Monomorphic proteins Ubiquitin, Galectin and Thermonuclease (PDB ID: 1UBQ, 2NN8 and 5KEE) **(B)** Experimentally known metamorphic proteins Kaib, RfaH and Lymphotactin (PDB ID: 5JYT and 2QKE, 2OUG and 2LCL, 1J8I and 2JP1). The experimentally solved structures are also depicted below each plot. The entropy score is plotted across residue number, the blue line plot is the entropy score for each fragment plotted corresponding to the residue number of the first residue, and the orange line plot is the rolling average of the entropy score with a window width of 18. The fold-switching regions show corresponding high entropy scores and mixed propensities, while the propensities of monomorphic proteins remain non-overlapping almost always.

In bioinformatics, substitution matrix-based scoring techniques are widely used. They are particularly helpful in quantifying the change or substitution of one element by another. We implemented a substitution matrix based scoring method in order to evaluate the ability of the fragments to adopt different secondary structure conformation. First, the secondary structure of the fragments in the original output from DSSP is taken. The original output from DSSP contains one of these 8 secondary structures: *α*-helix (H), 310 helix (G), π helix (I), poly-proline helix (P), extended strand (E),*β* bridge (B), turn (T), bend (S). A reference hit is set as the hit that has the most commonly occurring 7-mer secondary structure pattern among all the hits. If there is no majority, a random hit is assigned as the reference hit. Then every other hit is taken, pair-wise aligned with the reference hit and scored according to the following substitution matrix:

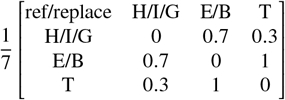

This way, as we align each fragment sequence hit with the reference hit and apply the substitution matrix on the alignment, we get a score. For a sliding window fragment, all pairwise alignment with the reference hit is taken and among them, if there are multiple hits from the same sequence cluster, these scores are weighted with the inverse of the total number of hits that we obtain from the same sequence cluster. We then take the weighted average of these scores from all pairwise alignments and assign it to that sliding window fragment. In this manner, we obtain a score for each fragment corresponding to all sliding windows.

We also sought to quantify the confidence in the calculated diversity, entropy and substitution scores from fragment-picking for a particular sliding window based on the number of hits we obtained for the corresponding fragment sequence. Ideally, there exists a true value of the propensity for the middle residue to be in a helical or sheet-like structure; the propensity that we arrive at is an estimate that gets better and better based on the number of fragment hits from different sequence clusters that we have available to us to make an estimation. From Hoeffding’s inequality [33], we know that the difference in the true value and the estimation varies with the sample size in an exponential manner. Hence, the uncertainty in the calculated values of diversity metrics is estimated in the form of an exponential function. Given a fragment with a large number of hits (say N), we can analyse how the propensities calculated from ‘n’ randomly selected fragments vary from the true propensity (n → N). This analysis helps in modelling the functional form of uncertainty. The final form for uncertainty is estimated by the function:

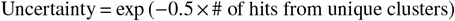

Rolling average over a range for functions that are irregular and discretely defined gives an output that is a smoothened representation of the initial function. They accumulate and average the variation of the discrete function across a window. We theorized that a protein sequence having a contiguous region of high entropy scores would imply that the region as a whole would have a higher chance of switching folds. Hence we chose to make a rolling average of the scores to elicit these contiguous regions. So, the score that is taken as input for classification is the maximum of the rolling average of the diversity, entropy and substitution scores. The rolling window width is a parameter that we are at liberty to choose. An optimal value for the window width is chosen using the training data. It is found that a window width of 8 and 18 gives the best separation of metamorphic and monomorphic scores in the training data (SI: Table S3). Hence a step function, as shown below, is used for the rolling average window:

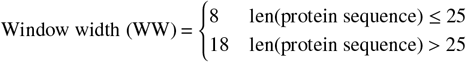

The final score that is used as the input vector for the classification model is all of the aforementioned scores together. After the rolling average window width is decided based on the length of the protein sequence, the maximum value of the rolling average of diversity, entropy and substitution scores is calculated. The mean value of uncertainty across all residues is also found and the four scores together are used as the feature space for the classification model, as shown below:

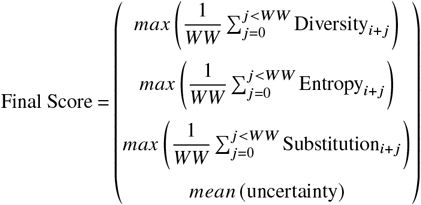

We noticed that when working with residues other than the middle residue in the 7-mer, similar levels of accuracy were achieved. The accuracies when using all the different residues (except the terminal residues) were within one standard deviation of each other. Among these, the model working with the third residue gave the highest accuracy, higher than that of the middle residue. (SI: Table S4). Moving forward, we decided to work with the third residue for the classification model.

In the formula for the final score, i ranges from 3 to L-7+3, where L is the length of the protein sequence. In the case that for a particular fragment, no hits are found, we set the value of the diversity metric scores to a minimum. Therefore, diversity is set to 1, entropy and substitution are set to 0. Uncertainty, in that case, achieves the maximum value of 1.0. This is done to reduce the false-positivity rate of the model in the case where we have no data to fix the propensity of the 3rd residue of the fragment and, therefore, the diversity of the fragment.

### 2.4 Prediction Model

A final prediction model is constructed using the aforementioned features. The training data is randomly split according to a 6-fold cross-validation scheme. Upon performing a 6-fold cross-validation, an SVM model with a polynomial kernel of degree 2 is found to be the model that has the highest average validation accuracy while also maintaining a lower model complexity. As a result, a quadratic SVM is selected to provide predictions on data that the model has never encountered before. Figure 3 shows the decision boundary that is finally obtained after optimizing the support vector machine using a Bayesian optimization scheme for the SVM hyperparameters. Table 1 shows the diversity metrics for the known and experimentally verified metamorphic proteins and a few monomorphic proteins.

**Table 1.**
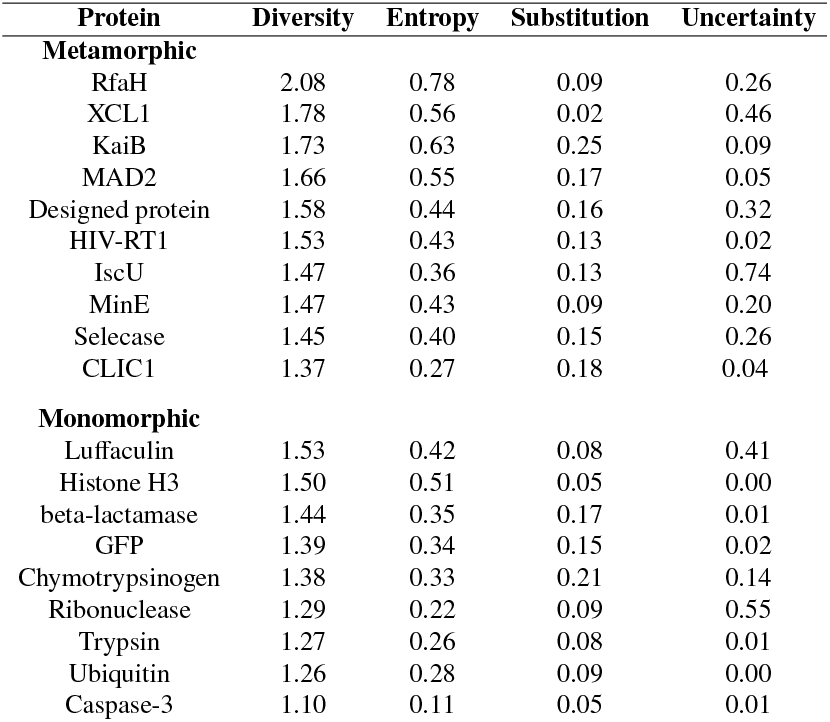
The diversity metrics of known metamorphic proteins and proteins that are highly likely to be monomorphic are ordered according to decreasing diversity scores.

**Figure 3.**
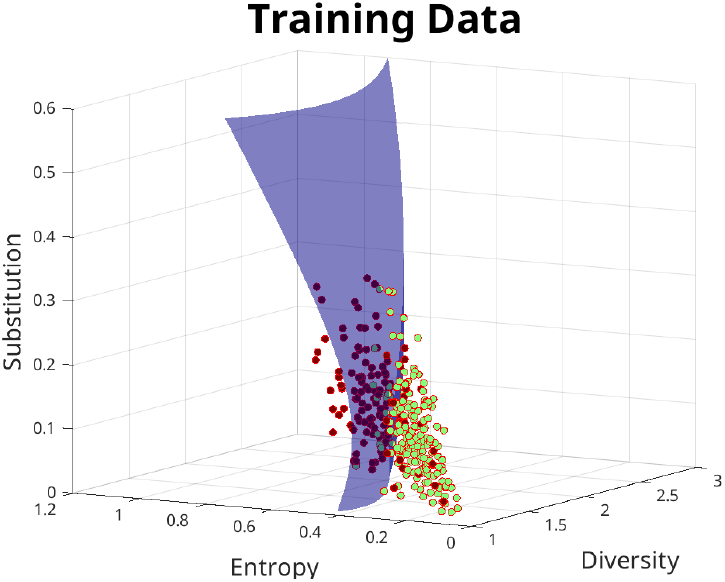
The training dataset with 3/4 features. The training dataset of known fold-switching proteins and highly likely monomorphic proteins are plotted in the feature space of the variables diversity score, entropy score and substitution score. The points in red depict fold-switching proteins and the points in green depict monomorphic proteins. The decision boundary shows the optimized SVM model output when a quadratic kernel is used.

Figs. S3(a-f) in SI show how the decision boundary varies as the model is re-trained across each of the 6 partitions. The corresponding validation confusion matrix is also shown in these figures. The standard deviations shown in Table 2 also give an idea of how much variation is present across the different folds used for training. Fig. S3(g) shows the ROC (Receiver-operating characteristic) curve for the final optimized SVM model. Although Figure 3 shows a 3D hyper-surface as the decision boundary, the actual decision boundary is a 4D hyper-surface corresponding to the feature space of the 4 scores used. The figure here is plotted for visualization purposes and to show the segregation of metamorphic and monomorphic proteins in the 3 feature space dimension. The 4 features are plotted in a parallel coordinates plot to see the mutual variation in all 4 coordinates with reference to each other (see Fig. S3(h) in SI). The final prediction model uses all 4 of the features to make new predictions.

**Table 2.** The validation accuracy measures for Morpheus and the method by Liwang’s group averaged across the cross-validation partitions are mentioned. The numbers in the parentheses are the standard deviation for these values calculated across the cross-validation partitions. The values for Chen et. al. directly taken from the values mentioned in the publication; precision and F1-score were inferred from the other values, hence we could not provide a standard deviation value.

| Measure | Morpheus | Chen et. al |
| --- | --- | --- |
| Sensitivity | 0.79 (0.08) | 0.63 (0.09) |
| Specificity | 0.90 (0.06) | 0.78 (0.06) |
| Precision | 0.89 (0.08) | 0.81 |
| Accuracy | 84.7% (6.1) | 69.8% (1.5) |
| F1 Score | 0.83 (0.08) | 0.71 |
| MCC | 0.698 (0.12) | 0.355 (0.104) |

### 2.5 Evaluation of Performance

The model performance is evaluated using different measures of accuracy. These measures include Accuracy (ACC), Sensitivity (Sn), Specificity (Sp), Precision, Matthew’s Correlation coefficient (MCC) and F1-score, which are commonly used measures for machine learning algorithms. They are defined as follows:

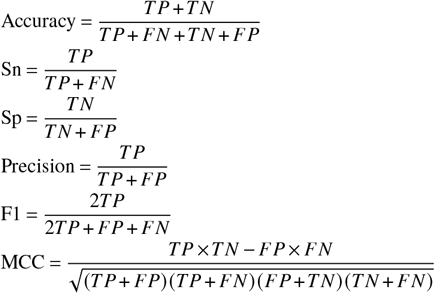

where TP, FP, TN, and FN denote the number of true positives, false positives, true negatives, and false negatives, respectively. Different classification models, such as Support Vector Machines (SVM), k-nearest neighbour (KNN), Linear Discriminant Analysis (LDA), Decision Tree, etc., are evaluated on the training data. Finally, one is picked to make predictions on data the model has not encountered while training. This decision is based on validation accuracy measures and lower model complexity. For example, although a fine Gaussian SVM gives high validation accuracy, the model complexity is quite high due to the fine nature of the kernel. Hence, to reduce the chances of overfitting, a model with lower complexity is preferred. An SVM model with a polynomial kernel of the second degree is selected as the prediction model of the classification models that are evaluated.

### 2.6 Work Flow of the Algorithm

The workflow of the algorithm is depicted in the schematic Fig. 4. The flow of information starts from the input query sequence, which is then used to perform fragment sequence search on the database, and then scored and fed into the SVM model. A binary output is produced, indicating whether the prediction for the query protein is metamorphic or monomorphic.

**Figure 4.**
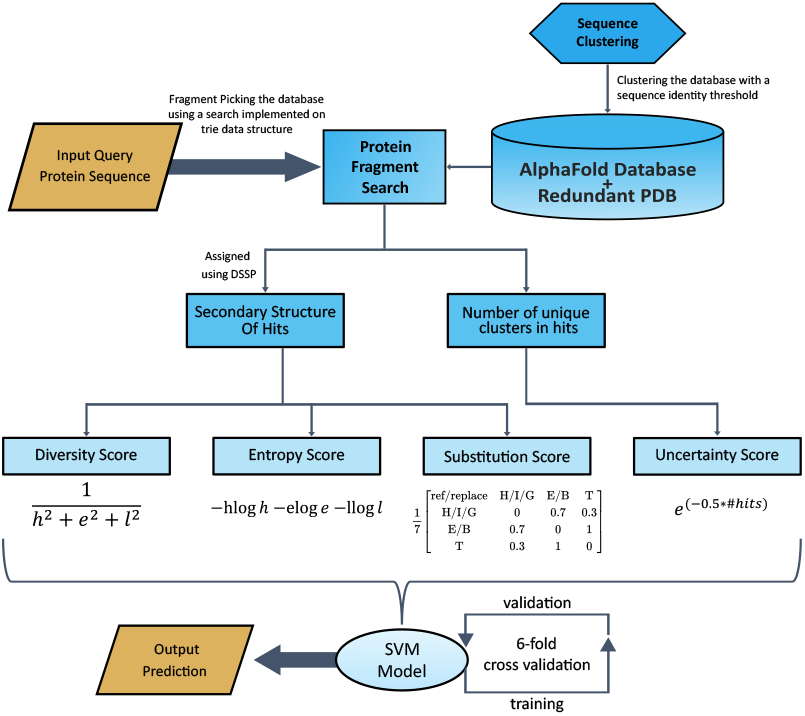
workflow diagram. Workflow is depicted starting with the fragment sequence search across the database. The hits are then scored for the diversity of secondary structures shown across different structures through 4 different scoring techniques. An SVM model with a quadratic kernel is employed with the scores as the features. The SVM model is trained and a 6-fold cross-validation scheme is applied. Finally, a binary output prediction is made using the trained SVM model.

## 3. Results and Discussion

Having constructed a classifier to predict the metamorphic behaviour in proteins using protein sequence information alone, we applied our algorithm to predict metamorphic proteins across 57 proteomes from different organisms. We also choose a few candidate metamorphic proteins from our predictions that are involved in crucial biological processes and are highly likely to switch folds based on a few criteria that are mentioned further in the results subsection.

### 3.1 Fragment structure based predictions outperforms purely sequence based algorithm

Our Model can predict metamorphic proteins with an average cross-validation accuracy of 84.7% and Matthew’s correlation coefficient of 0.698. Our method outperforms existing sequence-based metamorphic protein predictors while also maintaining a low false-positivity rate of 9.6%. Table 2 depicts the different measures of accuracy for the quadratic SVM model and existing sequence-based predictor of metamorphic proteins. Such measures of accuracy open up the exciting possibility of predicting metamorphic proteins at the proteome level.

Furthermore, plotting the diversity metrics and the secondary structure propensities at each residue allows us to gauge the approximate region that undergoes the fold switch. For example, in figure 2, residues around 51-100 of the known metamorphic protein KaiB show a contiguous region with a high entropy score. These residues overlap well with the experimentally shown region of fold-switching. This level of residue-wise information is expected to be useful when experiments are performed on candidate metamorphic proteins. Additionally, the nature of the fold-switch that may happen in the putative hits can also be inferred by analysing the secondary structure propensity plot of the candidate protein sequence.

### 3.2 Predictions across proteomes reveal varying widespread metamorphicity in proteins

As we now have an algorithm that is able to predict the metamorphic behaviour in proteins with high accuracy, we sought to make predictions on the proteomes of many different organisms. We have run our model on 57 different proteomes chosen from all kingdoms of life, from unicellular prokaryotes to multicellular eukaryotes. Analysing different proteomes would have distinct advantages; for example, analysing an archaean species would be beneficial, considering that we would be working with a proteome of smaller size and not have to worry about post-translational modification. We further noticed that viral proteins were more frequently populated in the training dataset of known fold-switching proteins, so we decided to make predictions with our algorithm on a consolidated database of viral proteins. All of the data for the 57 proteomes obtained from our predictions are made publicy available on the following Zenodo repository (https://zenodo.org/records/14837336). Furthermore, we offer a web server to execute the algorithm for proteins that are not included in our local runs (Morpheus).

Figure 5 shows the diversity metrics for all the proteins from the M. Tuberculosis, Yeast and Human proteomes. Fig. S4 in SI also display the proteome level data for a couple of other proteomes. The full dataset for the 57 proteomes is provided in the Zenodo server as an excel sheet for each proteome. Once we plot the diversity metrics for all the proteins, overlaying the decision boundary helps us make binary predictions of the metamorphic behaviour of those proteins along with a confidence score based on how far away from the decision boundary the points lie. We also notice that of the proteins that are predicted metamorphic by our algorithm, a good number of them are intrinsically disordered proteins (IDP). This is expected as our algorithm quantifies the diversity in secondary structures adopted by a particular fragment in the query protein with the help of a structural database. It so happens that IDPs that occupy a highly varying secondary structure across the different deposited structures are reported as containing a high diversity of secondary structures. When analysing the final prediction set, each protein can be labelled with the percentage of disordered residues that it contains with the help of existing tools for predicting disorder in proteins. As many of these tools achieve a high degree of accuracy, if needed, looking at a consensus on disorder prediction of a few different tools would help filter the dataset cleanly for the metamorphic sets.

**Figure 5.**
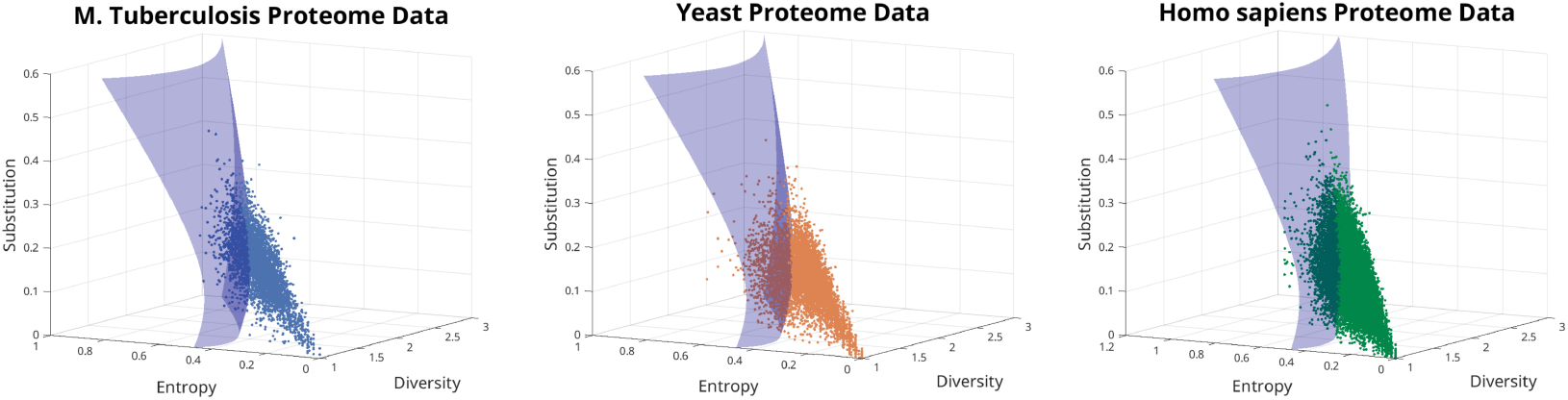
Distribution of proteins from 3 proteomes on the feature space overlayed with the decision boundary enabling new predictions of metamorphic proteins. 3995, 6060, and 20634 proteins from tuberculosis, yeast and the human proteome, respectively, are plotted as a scatter plot in the feature space of variables diversity score, entropy score and Substitution score. The SVM decision boundary is also plotted along with the data points. 831 (20.8%), 1289 (21.27%), and 4949 (23.98%) proteins from tuberculosis, yeast and the human proteome, respectively, are predicted metamorphic by our algorithm.

Figure 6 shows what fraction of the proteomes are predicted metamorphic by our algorithm. These numbers represent the entire prediction, including the intrinsically disordered proteins. To provide some distinctions, the bar plots are plotted along with a colour intensity gradient, which depicts the distribution of disorder among the proteins that are predicted to be metamorphic. As the field of metamorphic proteins is still in its infancy and there is quite a lot to uncover regarding their fold-switching mechanisms; the features that exactly define a metamorphic protein are still shaping up with ongoing research. Therefore, we decided to keep those proteins that are predicted to be intrinsically disordered in the final prediction set. We believe the conformational plasticity in proteins is a spectrum ranging from entirely monomorphic proteins on one end, which show the lowest conformational heterogeneity; to intrinsically disordered proteins, which show the highest conformational heterogeneity and lack a stable secondary structure most of the time. In the middle lies a set of proteins, including metamorphic, fold-switching, molten globules and multi-domain proteins with disordered regions. The distinction between metamorphic, fold-switching and proteins with disordered regions is not a clear distinction, yet. Hence, discarding proteins containing regions of disorder from the prediction set would be introducing bias and disregarding the continuum of conformational heterogeneity.

**Figure 6.**
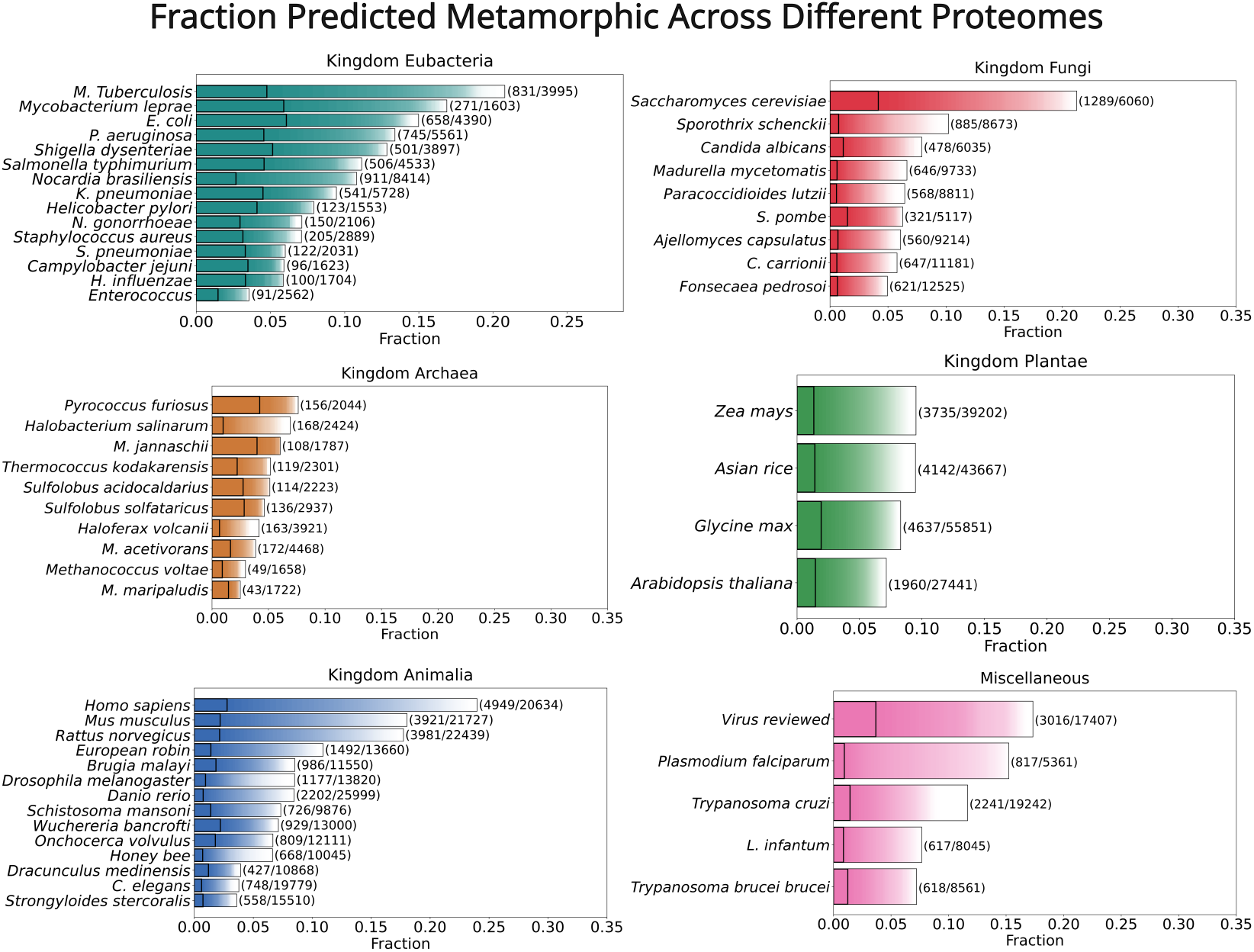
The distribution of metamorphic proteins predicted by Morpheus across different proteomes. The fraction of proteins predicted metamorphic across proteomes of different species. Among those predicted metamorphic, the distribution of disordered proteins is shown through the colour intensity gradient. The disorder prediction is performed on each protein that is predicted metamorphic, and the fraction of residues that are predicted to be disordered is calculated. In the colour intensity gradient, *α* = 1 (full intensity) represents completely ordered proteins, and *α* = 0 (white) represents completely disordered. The vertical black line denotes the fraction of proteins that have ≤ 1% residues predicted to be disordered. The plots are segregated into subplots based on the different kingdoms of life, and the final subplot titled miscellaneous contains viral species (consolidated database of manually reviewed proteins from viral proteomes) and from other kingdoms such as Excavata and Protista.

After having made the prediction across different proteomes, we can analyse the results. The set of proteins that are predicted to be metamorphic with high confidence is a promising source for recruiting potential candidates that can be further tested for metamorphic behaviour through experimentation. In the following section, we provide a filtered set of proteins that are picked from the prediction set, keeping a certain set of criteria previously mentioned in mind.

### 3.3 Shortlisted candidate proteins for possible further investigation

In the search for strong candidate metamorphic proteins, we had the following criteria in mind to filter the prediction dataset: **(a)** The protein sequence should be predicted metamorphic according to Morpheus with high confidence, **(b)** the protein should have a range of residues where the helix and sheet propensity values are both non-zero for a contiguous region indicating the potential ability to fold-switch, and **(c)** the fraction of residues predicted to be disordered should be less than 10% and not in the region of fold-switch predicted by Morpheus. The threshold 10% is placed to account for the disorder region at the C or N terminus, if present. The disorder prediction is performed by IUPRED2A [34]. Based on these criteria, we select a few of these proteins and present them here in this section.

Table 3 lists a few proteins that are predicted to be metamorphic and are good candidates to be considered for further experimentation. The list contains two protein sequences, CXCL11 and MAD2B, which are related to already known metamorphic proteins that are lymphotactin and MAD2A. The sequences of CXCL11 and MAD2B are not similar to their metamorphic counterparts. CXCL11 already has two solved structures deposited in the PDB solved via NMR and Electron microscopy. Although the two structures have different secondary structures, the fold-switching has not yet been studied. The NMR structure gives us clues of CXCL11 being a metamorphic protein with temperature being the trigger [35]. So far, only one protein that is part of the circadian rhythm has been experimentally shown to be metamorphic in nature [11]. Since circadian rhythm is periodic and helps the organism in synchronizing with the time of the day, it would be quite an exciting find, if a new metamorphic protein joins the circadian rhythm. In our dataset, one of the promising protein predictions involved in the circadian rhythm is ‘Photolyase/cryptochrome alpha/beta domain-containing protein’ which is part of the plant proteome of Maize. In plants, cryptochrome acts as a Blue light receptor to entrain circadian rhythms. They also mediate a variety of light responses, such as the regulation of flowering and seedling growth. [36]. However, there are no solved structures for this particular protein in maize. Hence, this is a possible avenue for further research. Please see Fig. S5(a-j) where the diversity metrics are plotted for the selected few predictions listed in the paper. The plots contain calculated helix and sheet propensities, the entropy scores, the diversity scores and Substitution scores at each sliding window. The uncertainty score at each fragment is also taken into a rolling average and overlayed on the diversity score.

**Table 3.** List of potential candidates for metamorphic proteins. 6 of the 10 protein sequences marked with * do not have any experimentally solved structures deposited in the PDB as of when this manuscript was written.

| Uniprot ID | Organism | Protein Name | Length | Confidence | Uncertainty |
| --- | --- | --- | --- | --- | --- |
| O14625 | Homo Sapiens | CXCL11 | 94 | 6.48 | 0.52 |
| Q9Y2Y1 | Homo Sapiens | DNA-directed RNA polymerase III subunit RPC10 | 108 | 4.83 | 0.18 |
| Q4CMH7* | Trypanosoma cruzi | Guanine nucleotide-binding protein subunit beta-like protein | 102 | 4.80 | 0.44 |
| P26995 | P. aeruginosa | Transcriptional anti-activator ExsC | 145 | 2.82 | 0.43 |
| Q9UI95 | Homo Sapiens | Mitotic spindle assembly checkpoint protein MAD2B | 211 | 2.47 | 0.01 |
| A0A1D6P714* | Maize | Extensin-like protein | 129 | 2.27 | 0.61 |
| A0A804PFY2* | Maize | Photolyase/cryptochrome alpha/beta domain-containing protein | 162 | 1.86 | 0.45 |
| A0A3P7DY23* | Wuchereria bancrofti | S1 motif domain-containing protein | 258 | 1.23 | 0.39 |
| Q4DL46* | Trypanosoma cruzi | Copper-transporting ATPase-like protein, putative | 263 | 1.03 | 0.25 |
| B8A185* | Maize | Secreted protein | 112 | 0.72 | 0.34 |

## 4. Scope and Limitation

Conventionally, the two conformations of metamorphic proteins are experimentally validated by protein structure determination techniques like NMR and cryo-electron microscopy, or large changes in the secondary structure of proteins can be found through circular dichroism [37]. These methods can prove to be resource-intensive and time-consuming. To add to that, the trigger for fold-switching may not be known beforehand, making the detection of metamorphic proteins further difficult. Therefore, providing a way to systematically filter out monomorphic proteins and to find candidate metamorphic proteins only using the sequence information of the protein is highly useful. Our work is the first implementation of a proteome-wide metamorphic protein predictor using protein sequence information. Morpheus allows for the prediction of proteins whose structure may not be deposited in the PDB. Hence, our method is a step in the direction of expanding the known set of metamorphic proteins. Besides fulfilling a researcher’s curiosity, analysing the human proteome for metamorphic proteins can stand a chance at helping global healthcare and improving therapeutics by furthering the understanding of protein fold switching and the different physiological triggers involved within. Fold-switching proteins associated with a number of diseases have already been identified [38–41]. Broadening the reservoir of predicted metamorphic proteins help in paving ways for new research.

Although we are able to make high-throughput predictions of metamorphic behaviour in proteins to a good accuracy, our algorithm falls short at making sensitive predictions when the sequences differ by point mutations. For example, E48A in RfaH has been shown to disrupt the inter-domain contact and thus favour all beta conformation [9, 42]. So while analysing highly similar homologous sequences, the results obtained from our algorithm may be questionable. Further, since our algorithm takes into account the primary and secondary structure information of proteins, metamorphic proteins whose secondary structure remains intact, but there are rearrangements in tertiary contacts and hydrogen bonding would likely not be picked up by our model.

## Acknowledgments

AS would like to acknowledge Sridip Purui for carefully reading the manuscript and providing valuable comments. We acknowledge the RCSB team for providing optimized code to perform linear fragment-picking searches on large databases.

AS acknowledges the HPC facility “Beagle” that was set up from grants by the erstwhile IISc-DBT partnership program. AS thanks the DST for the National Supercomputing Mission grants (DST/NSM/R&D-HPC-Applications/2021/03.10, DST/NSM/R&D-HPC-Applications/Extension Grant/2023/27). AS also acknowledges the FIST program sponsored by the Department of Science and Technology, India that supports the MBU infrastructure. AS would also like to thank the Teams Science Grant from the DBT-Wellcome Trust India Alliance (Grant number: IA/TSG/21/1/600245). AS also thanks the DBT National Network Project (NNP) grant (BT/PR40323/BTIS/137/78/2023) and the Matrics grants (MTR/2023/001040) from the Science and Engineering Board (SERB), India.

## Author contributions

AS conceptualized the project. RA and AS designed the research. RA, VS and AS formalized the algorithm. VS and RA performed the research and analyzed the data. VS wrote all the codes, curated the fragment dataset and ran the algorithm at the proteome scale. HV worked on the web server. AS supervised the study. VS prepared the first draft of the paper with inputs from AS. All authors helped in polishing the draft.

## Data Availability

The code used for fragment-picking analysis is publicly available on Zenodo (https://zenodo.org/records/14837336). The Zenedo repository also contains the proteome-level prediction data for the 57 proteome. The algorithm is also hosted within the web server (http://mbu.iisc.ac.in/∼anand/morpheus).

## Supplementary Information

### 1 Supplementary Figures

#### 1.1 Figure S1: Schematic

Figure S1.a shows the fragment picking schematic with the cartoon representation of the fragments from different hits and their respective PDB ID/AlphaFold Accession numbers.

**Figure S1a:**
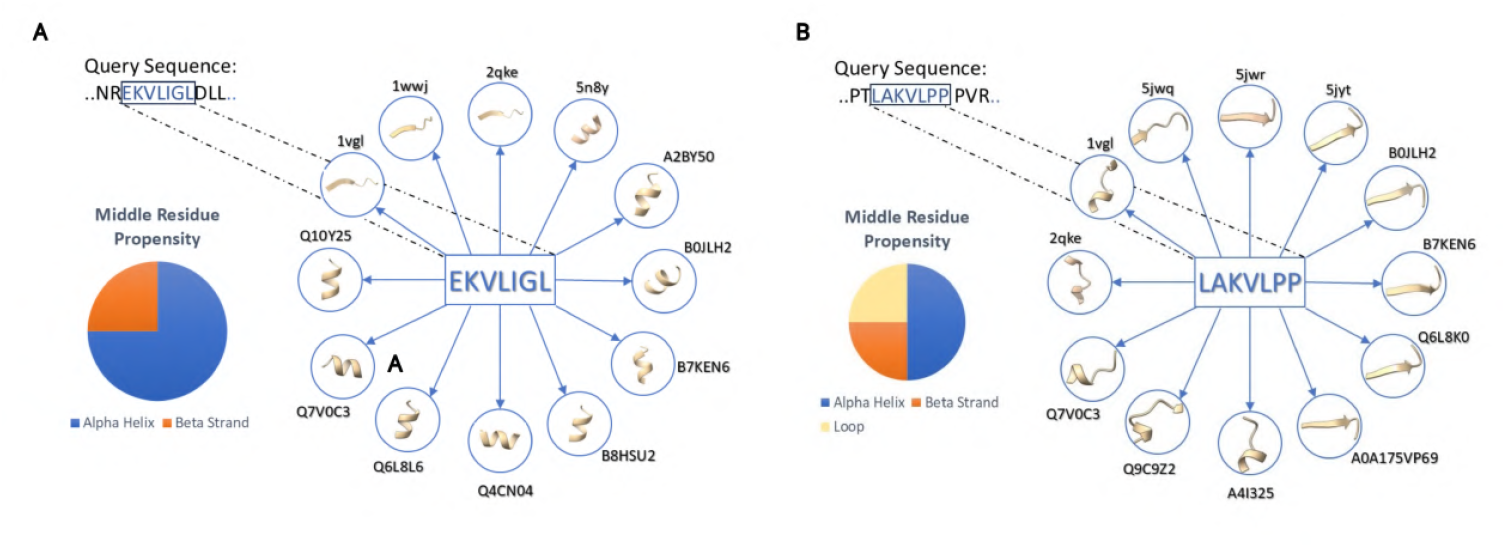
Schematic for fragment picking. The fragment sequences shown are part of the metamorphic protein KaiB **A** EKVLIGL and **B** LAKVLPP

#### 1.2 Figure S2: Diversity Metrics

The 10 figures in S2 show the diversity metrics (diversity, information entropy, substitution score and uncertainty) for 10 different experimentally known metamorphic proteins. They are RfaH, KaiB, IscU, Mad2, Lymphotactin, Selecase, MinE, CLIC1, HIV-RT1, and the designed metamorphic protein. The plots contain calculated helix and sheet propensities, the entropy scores, the diversity scores and Substitution scores at each sliding window. The uncertainty score at each fragment is also taken into a rolling average and overlayed on the diversity score. The experimentally solved structures for the two conformations are also displayed.

**Figure S2a:**
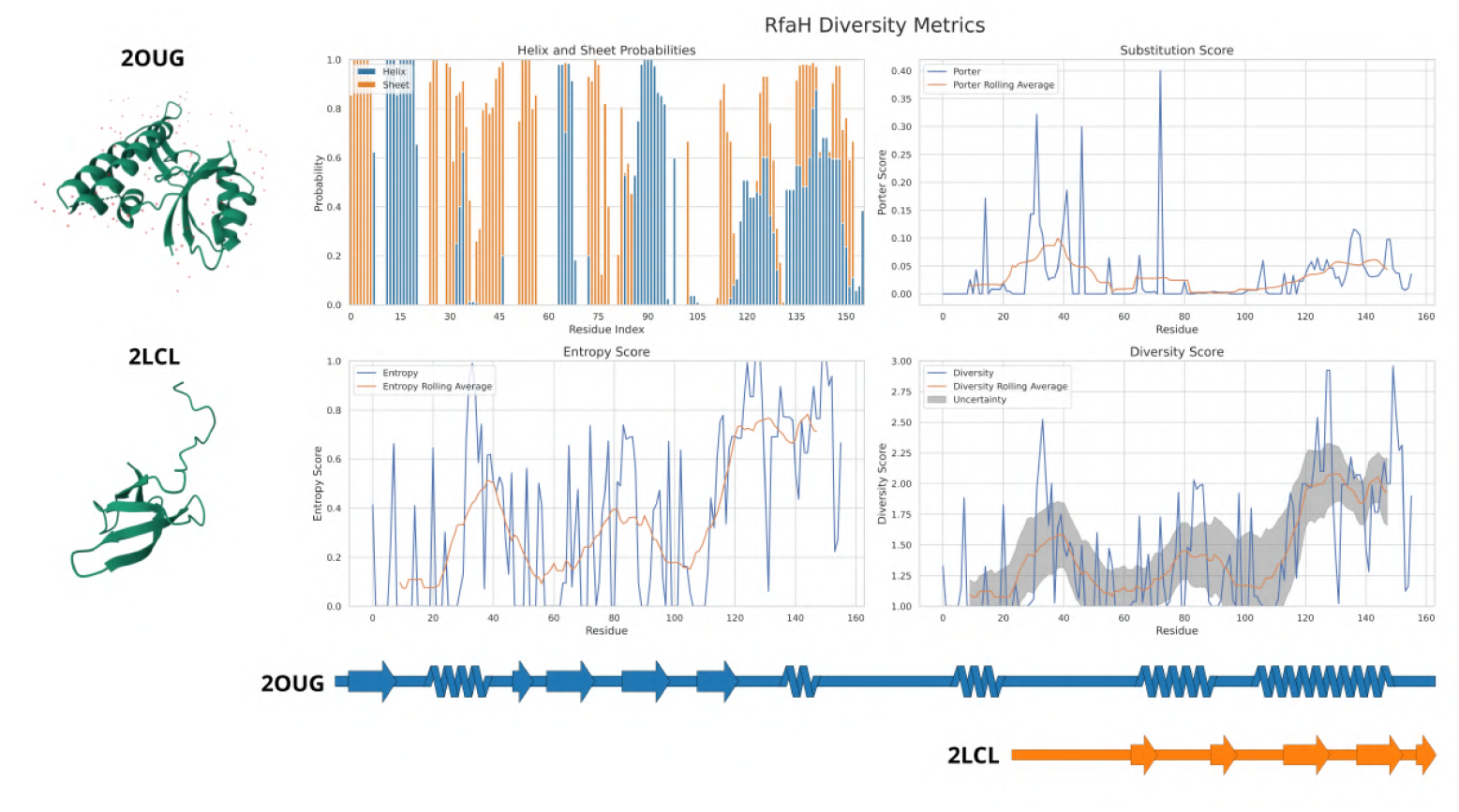
RfaH: Diversity metrics and secondary structure information for the metamorphic protein RfaH. The corresponding PDB IDs are 2OUG and 2LCL.

**Figure S2.b:**
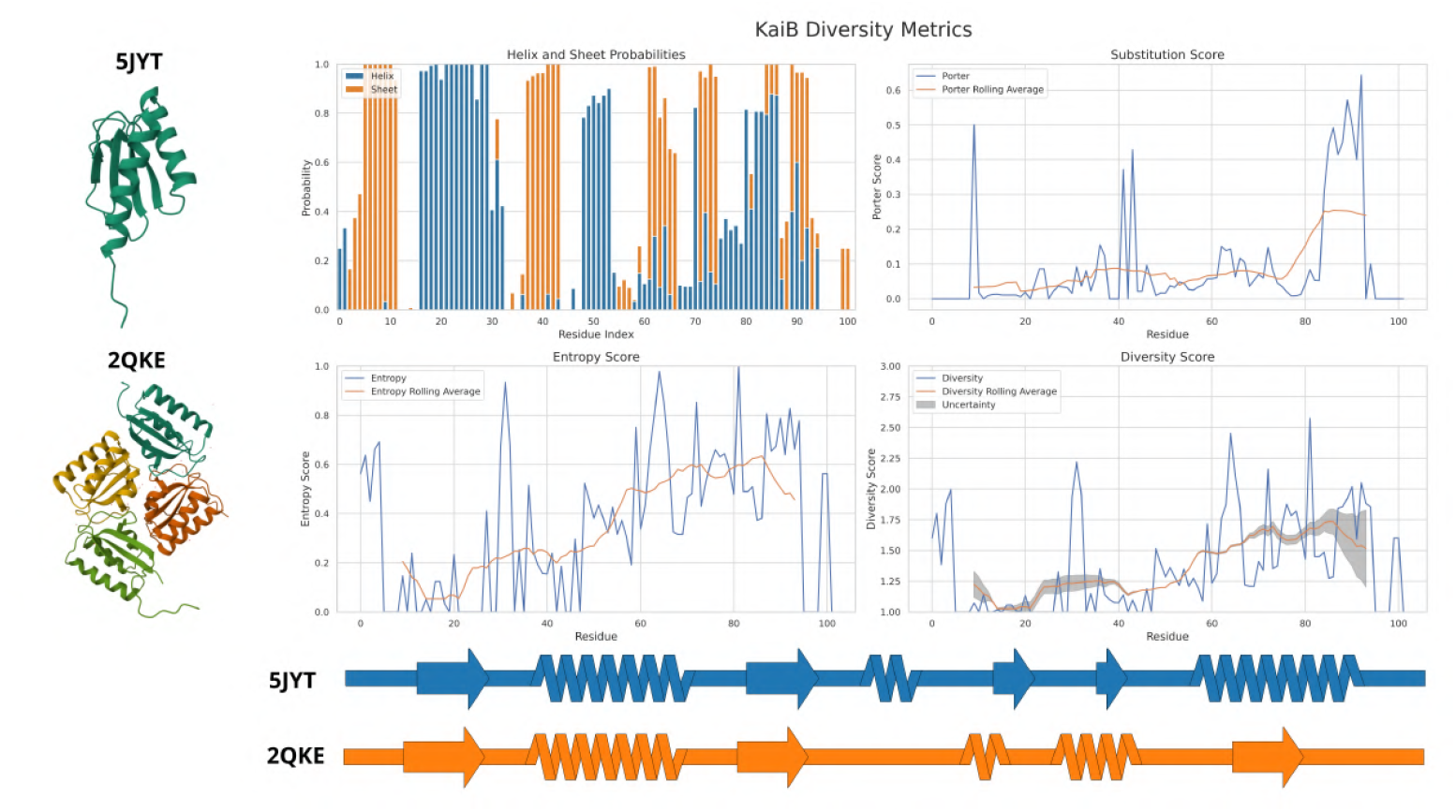
KaiB: Diversity metrics and secondary structure information for the metamorphic protein KaiB. The corresponding PDB IDs are 5JYT and 2QKE.

**Figure S2.c:**
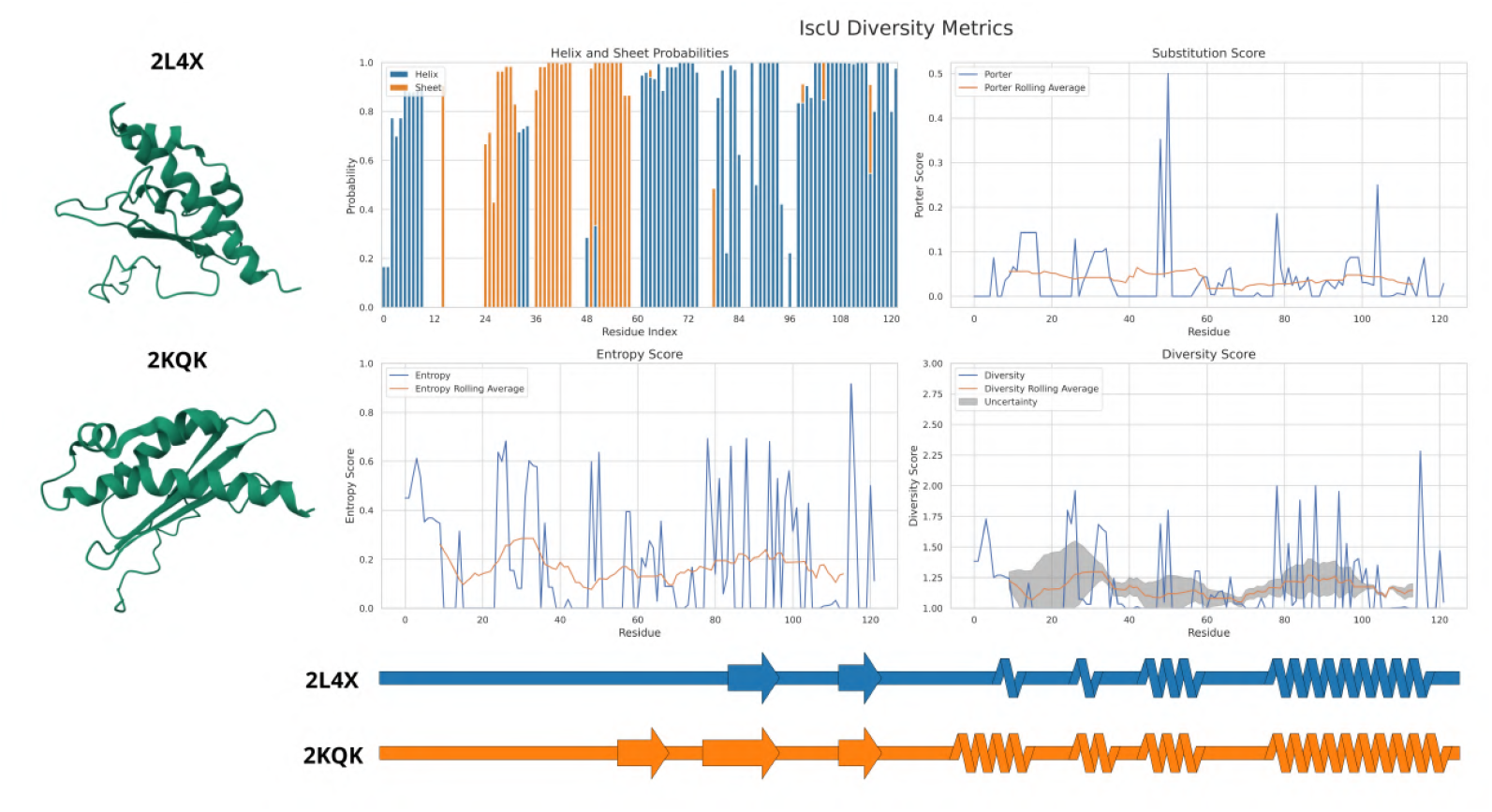
IscU: Diversity metrics and secondary structure information for the metamorphic protein IscU. The corresponding PDB IDs are 2L4X and 2KQK.

**Figure S2.d:**
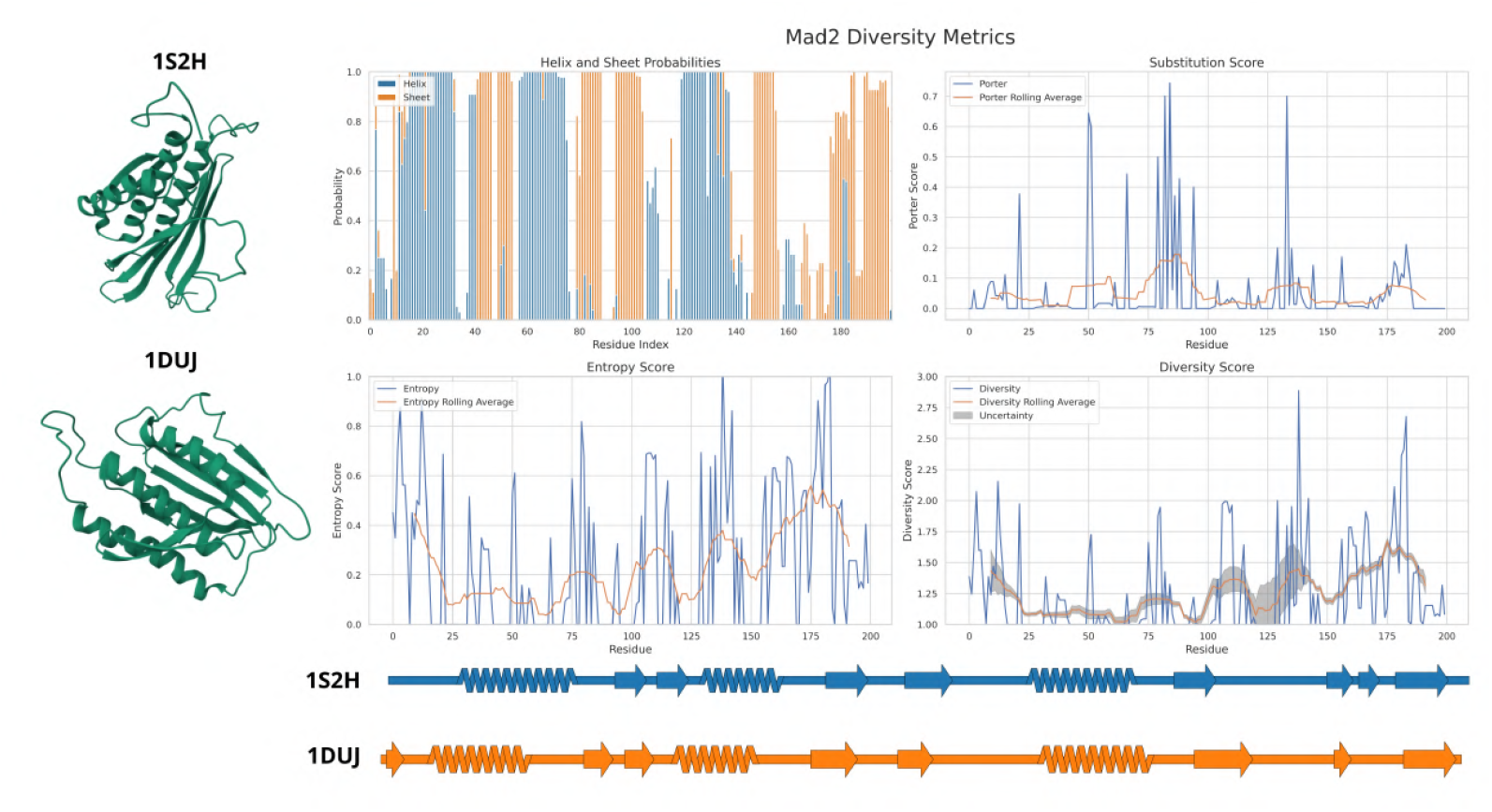
MAD2: Diversity metrics and secondary structure information for the metamorphic protein MAD2. The corresponding PDB IDs are 1S2H and 1DUJ.

**Figure S2.e:**
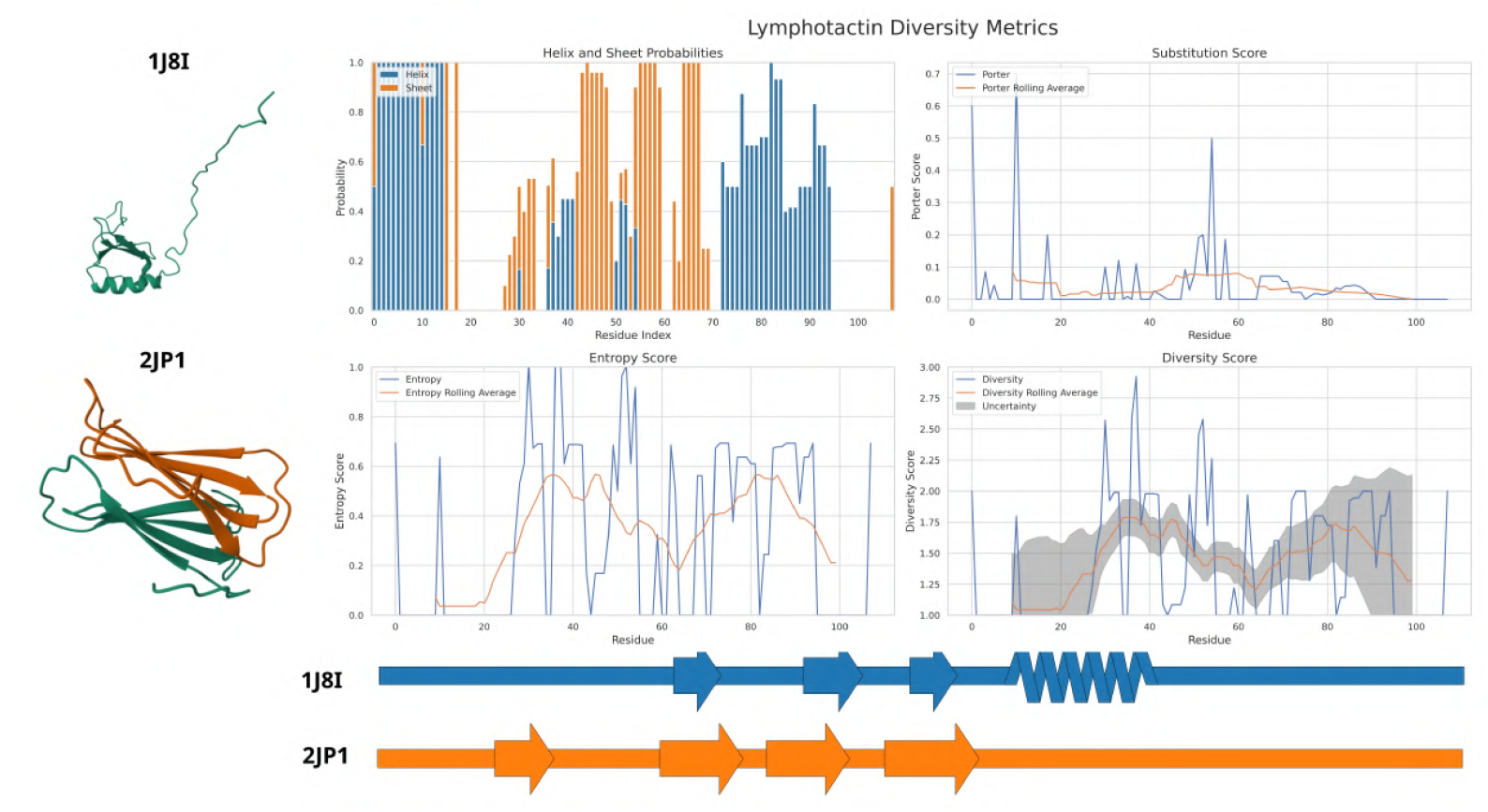
Lymphotactin: Diversity metrics and secondary structure information for the metamorphic protein Lymphotactin. The corresponding PDB IDs are 1J8I and 2JP1.

**Figure S2.f:**
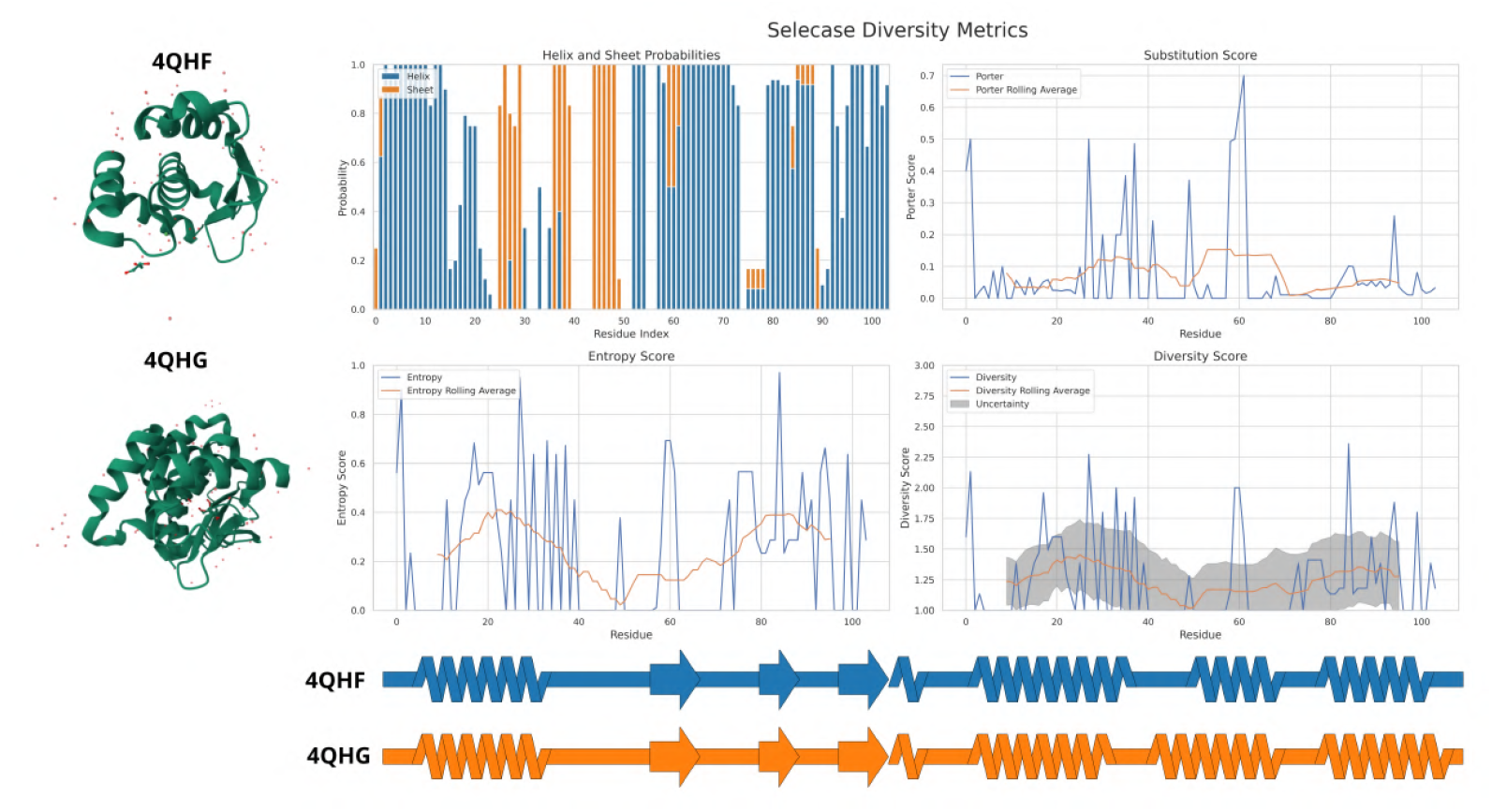
Selecase: Diversity metrics and secondary structure information for the metamorphic protein Selecase. The corresponding PDB IDs are 4QHF and 4QHG.

**Figure S2.g:**
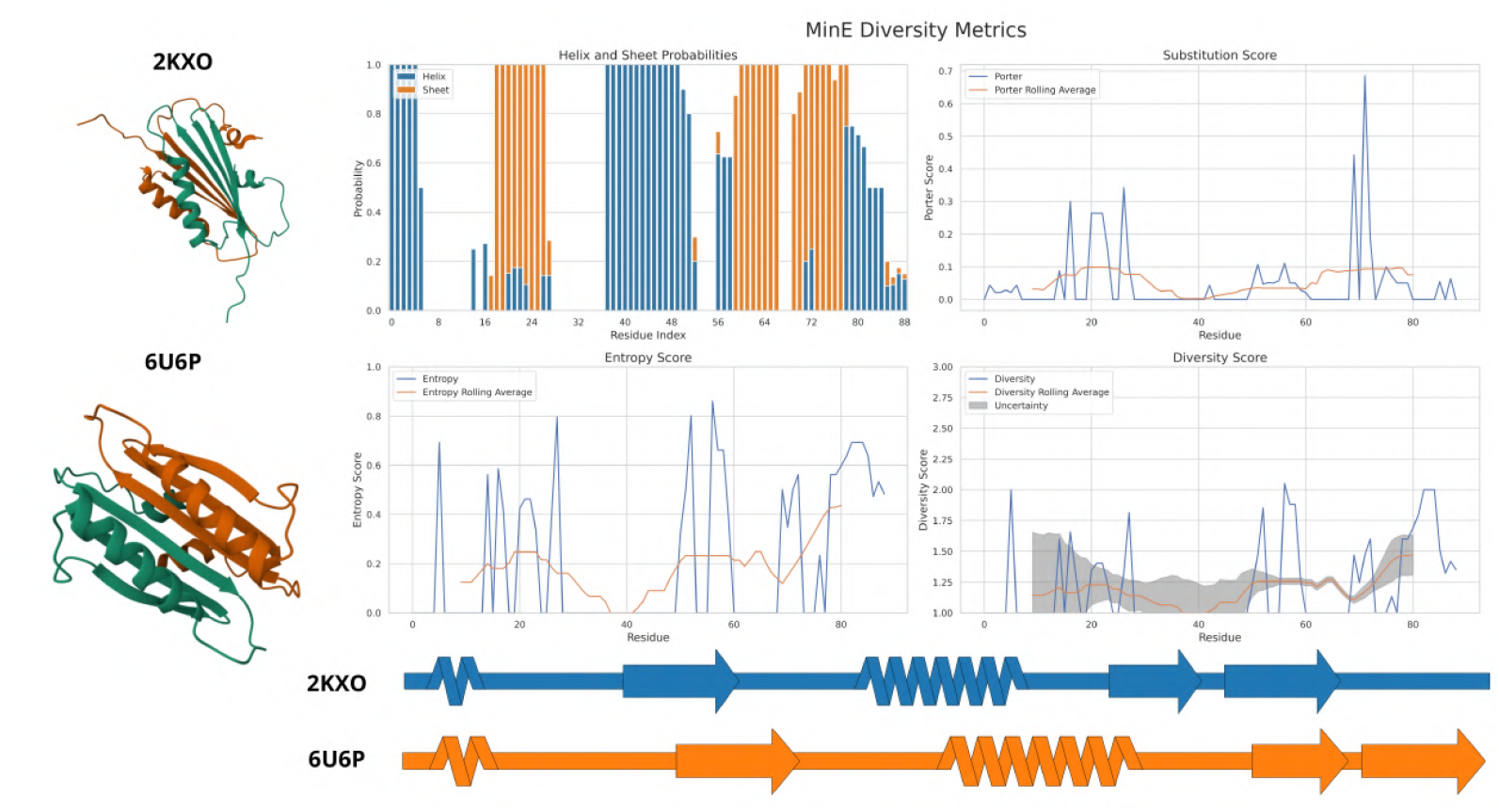
MinE: Diversity metrics and secondary structure information for the metamorphic protein MinE. The corresponding PDB IDs are 2KXO and 6U6P.

**Figure S2.h:**
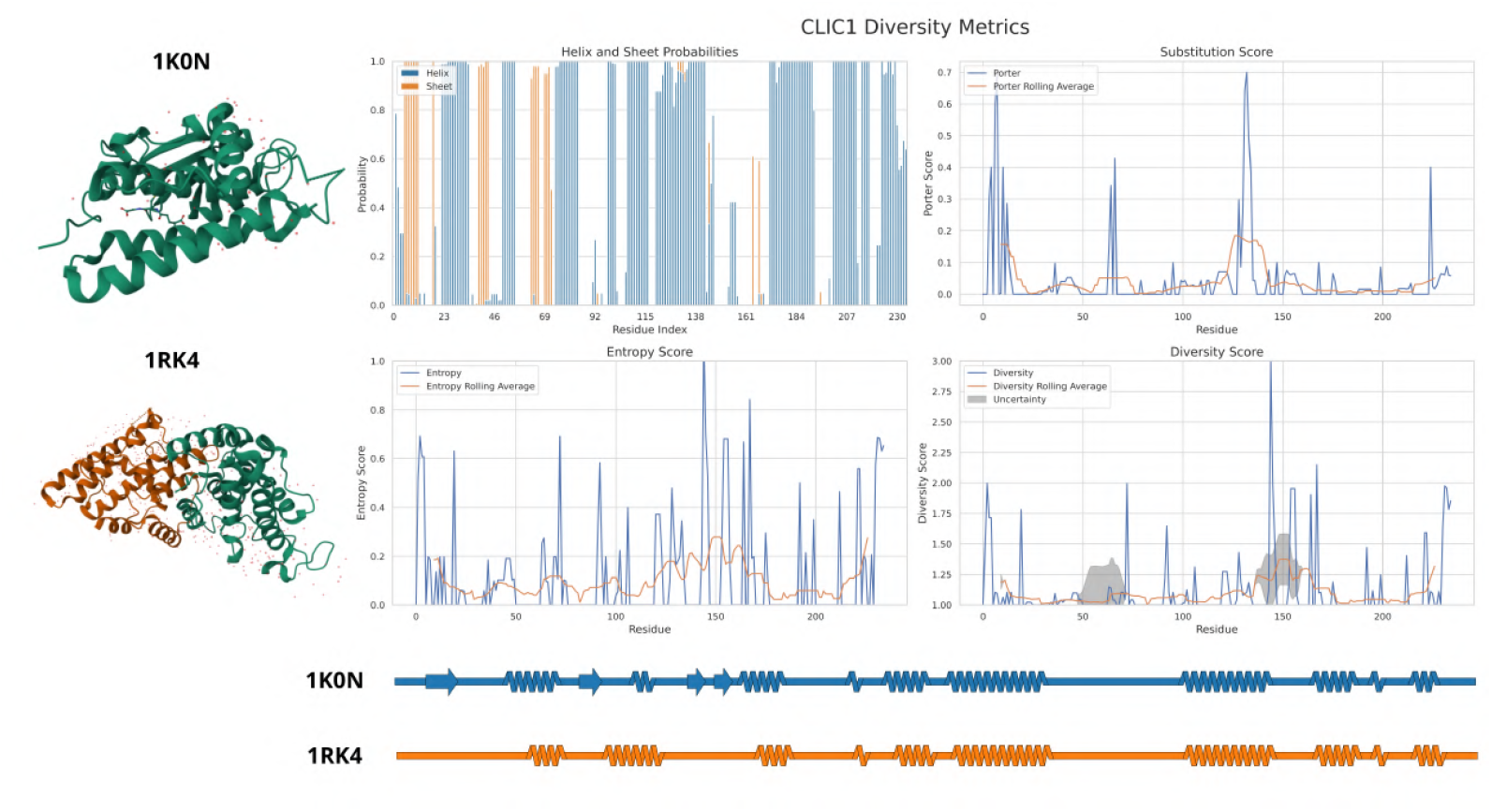
CLIC1: Diversity metrics and secondary structure information for the metamorphic protein CLIC1. The corresponding PDB IDs are 1K0N and 1RK4.

**Figure S2.i:**
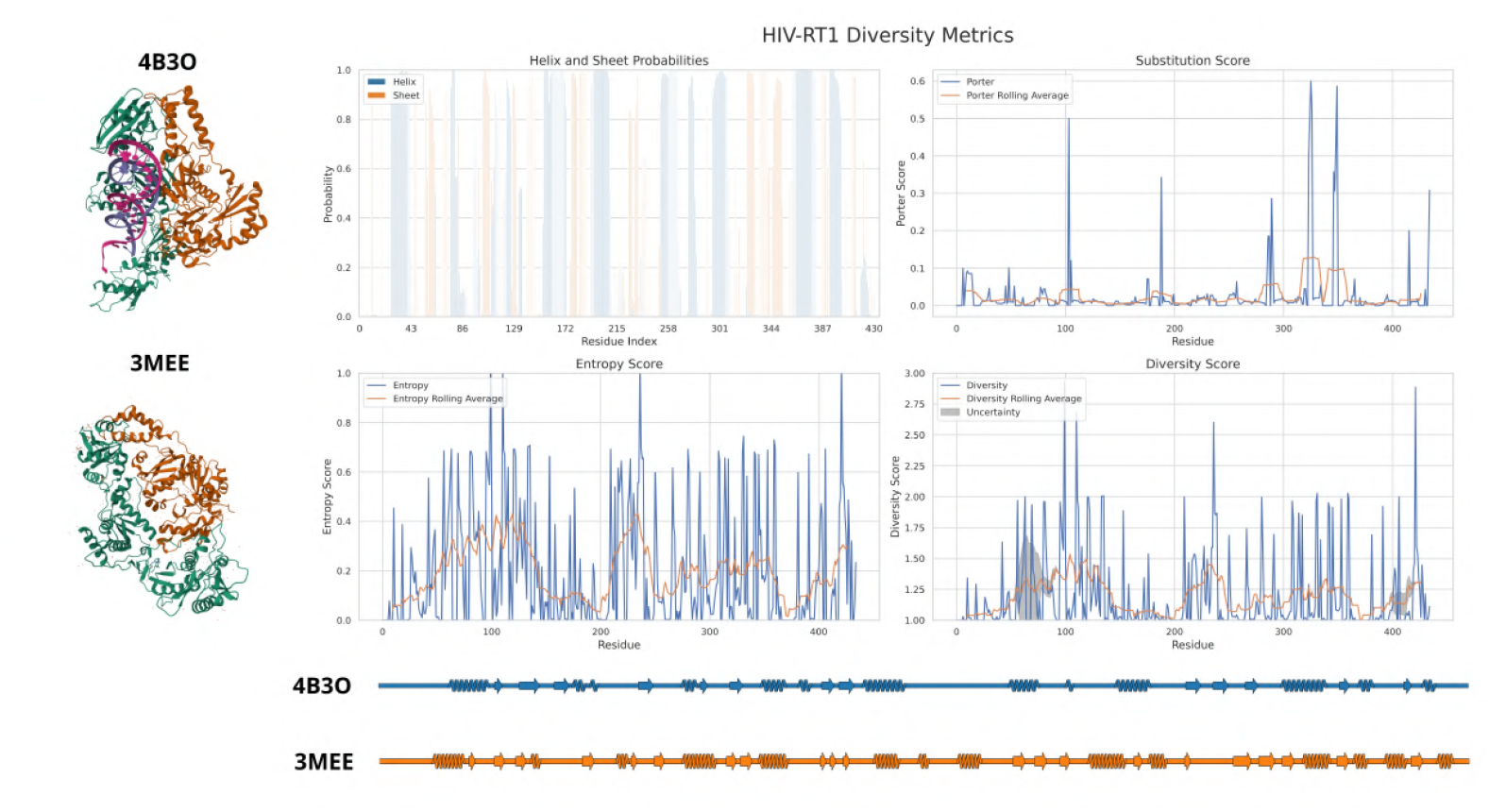
HIV-RT1: Diversity metrics and secondary structure information for the metamorphic protein HIV Reverse Transcriptase 1. The corresponding PDB IDs are 4B3O and 3MEE.

**Figure S2.j:**
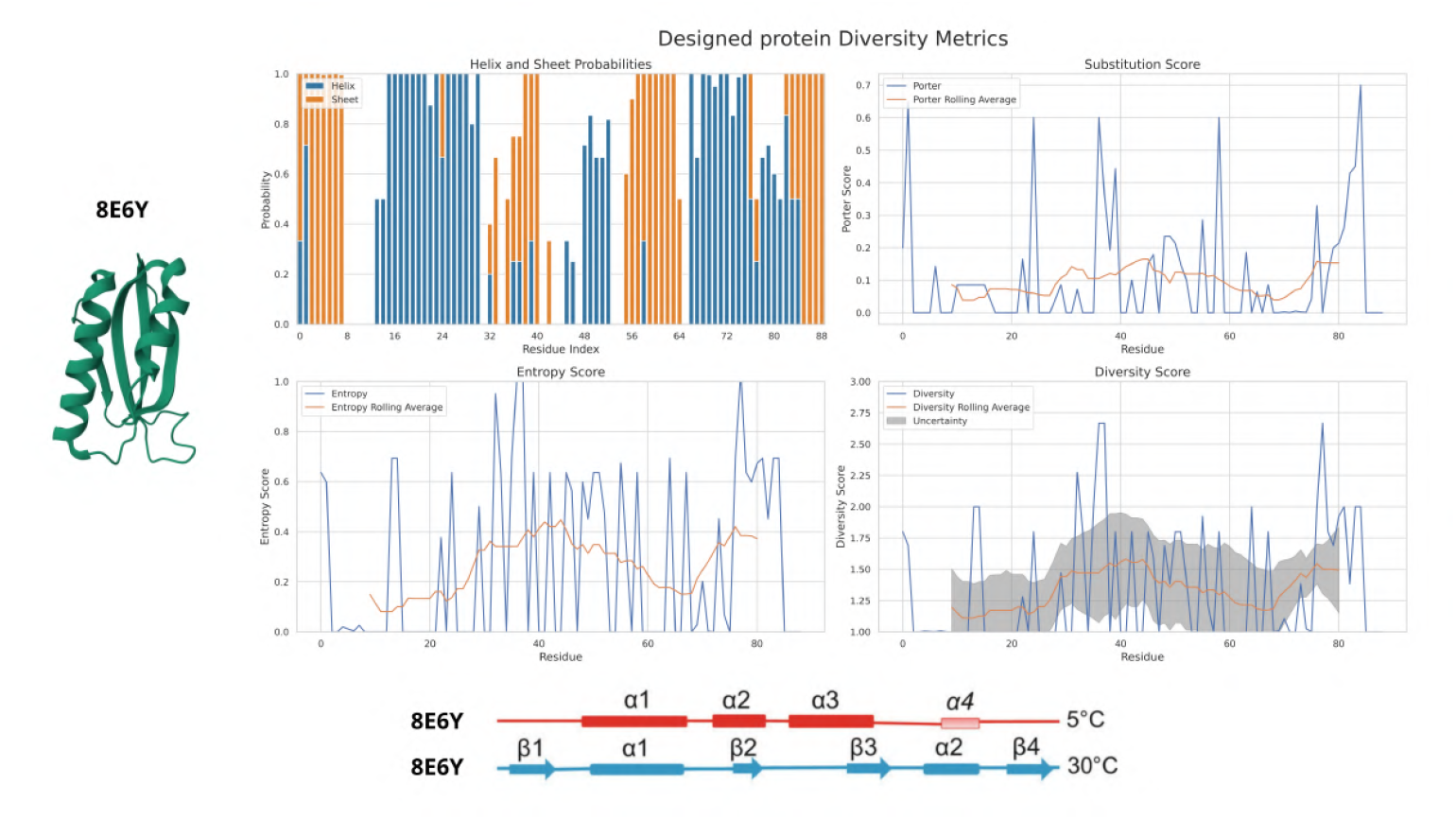
Designed Protein: Diversity metrics and secondary structure information for designed metamorphic protein. The corresponding PDB ID is 8E6Y.

#### 1.3 Figure S3: Cross-validation

Figures S3 show how the decision boundary changes when the quadratic SVM model is trained with the 6 different splits of 6 fold cross-validation. The corresponding validation confusion matrix is also shown. Figure S3.f shows the ROC curve and aggregate confusion matrix (all 6 folds) for the final trained SVM model. S3.h shows the parallel coordinate plot and *χ*^2^ scores from feature selection test for the four diversity metrics.

**Figure S3a:**
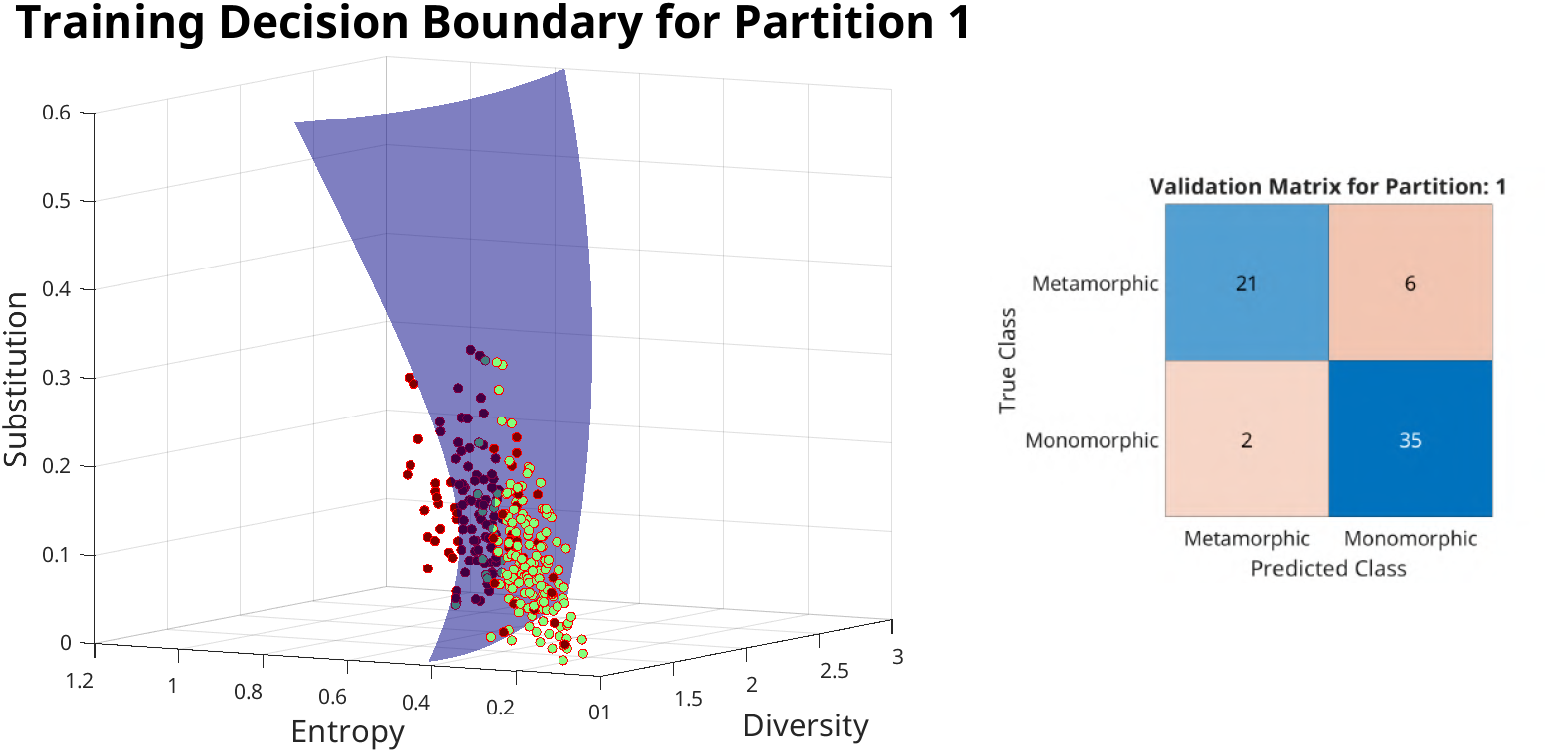
Decision boundary and the corresponding validation confusion matrix for the first training split partition

**Figure S3.b:**
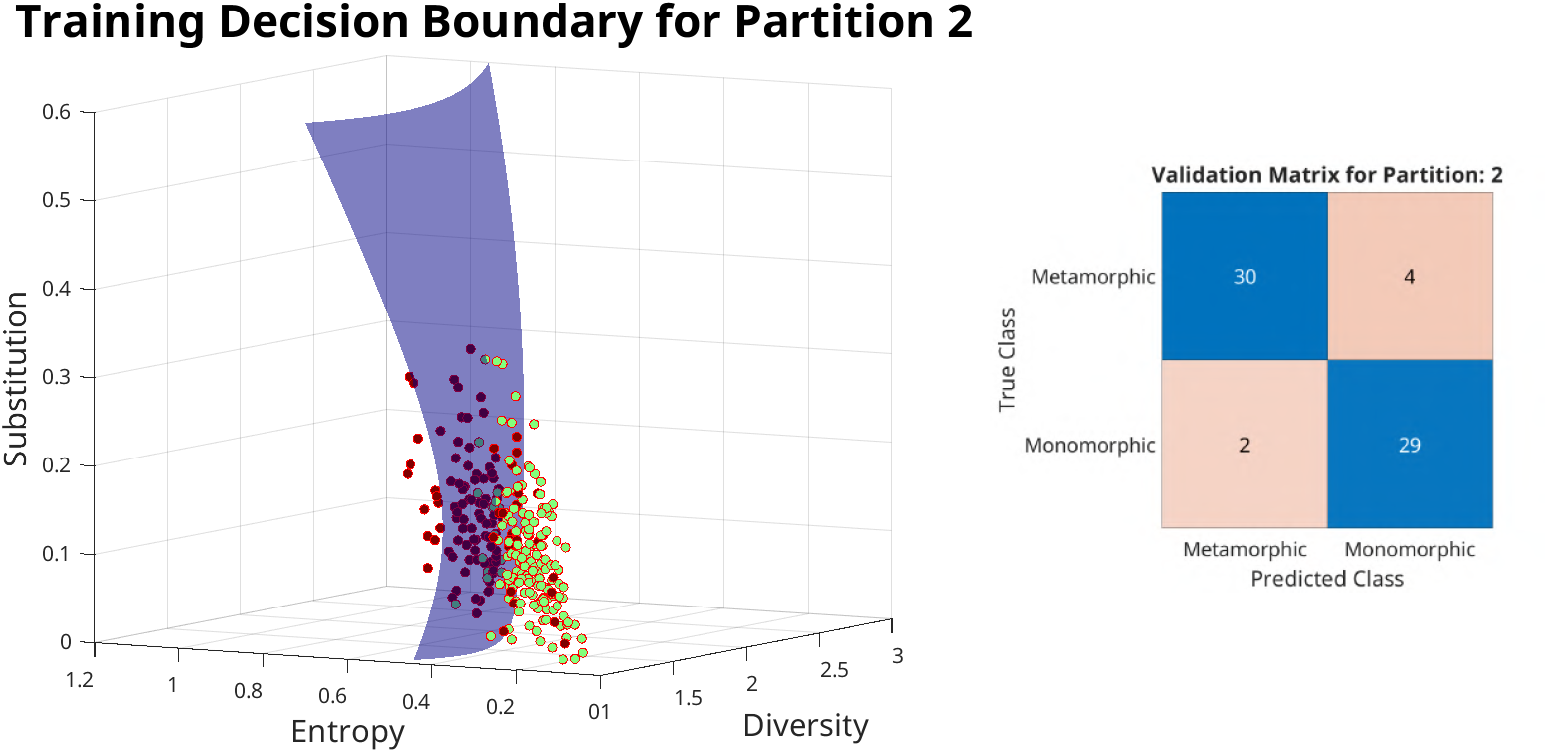
Decision boundary and the corresponding validation confusion matrix for the second training split partition

**Figure S3.c:**
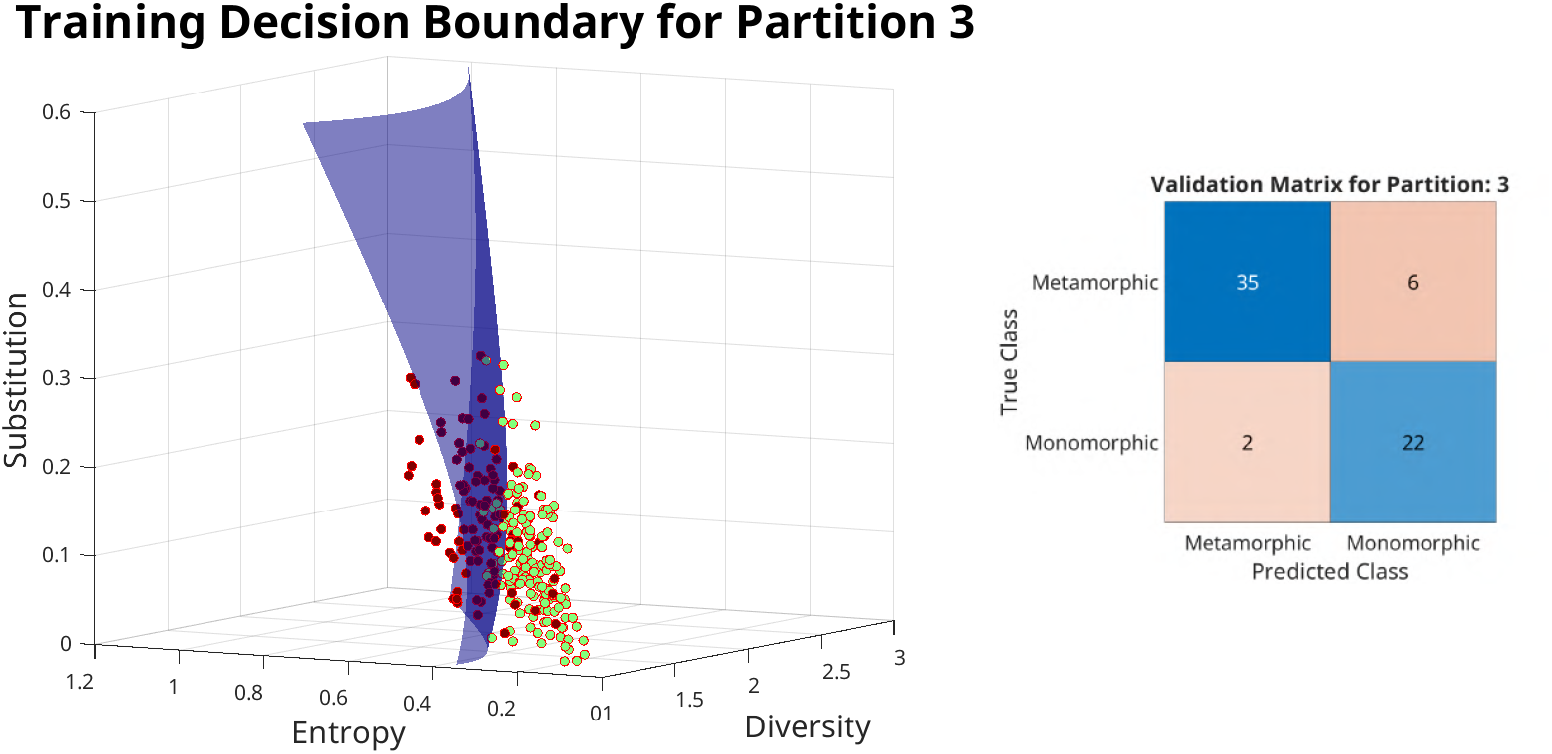
Decision boundary and the corresponding validation confusion matrix for the third training split partition

**Figure S3.d:**
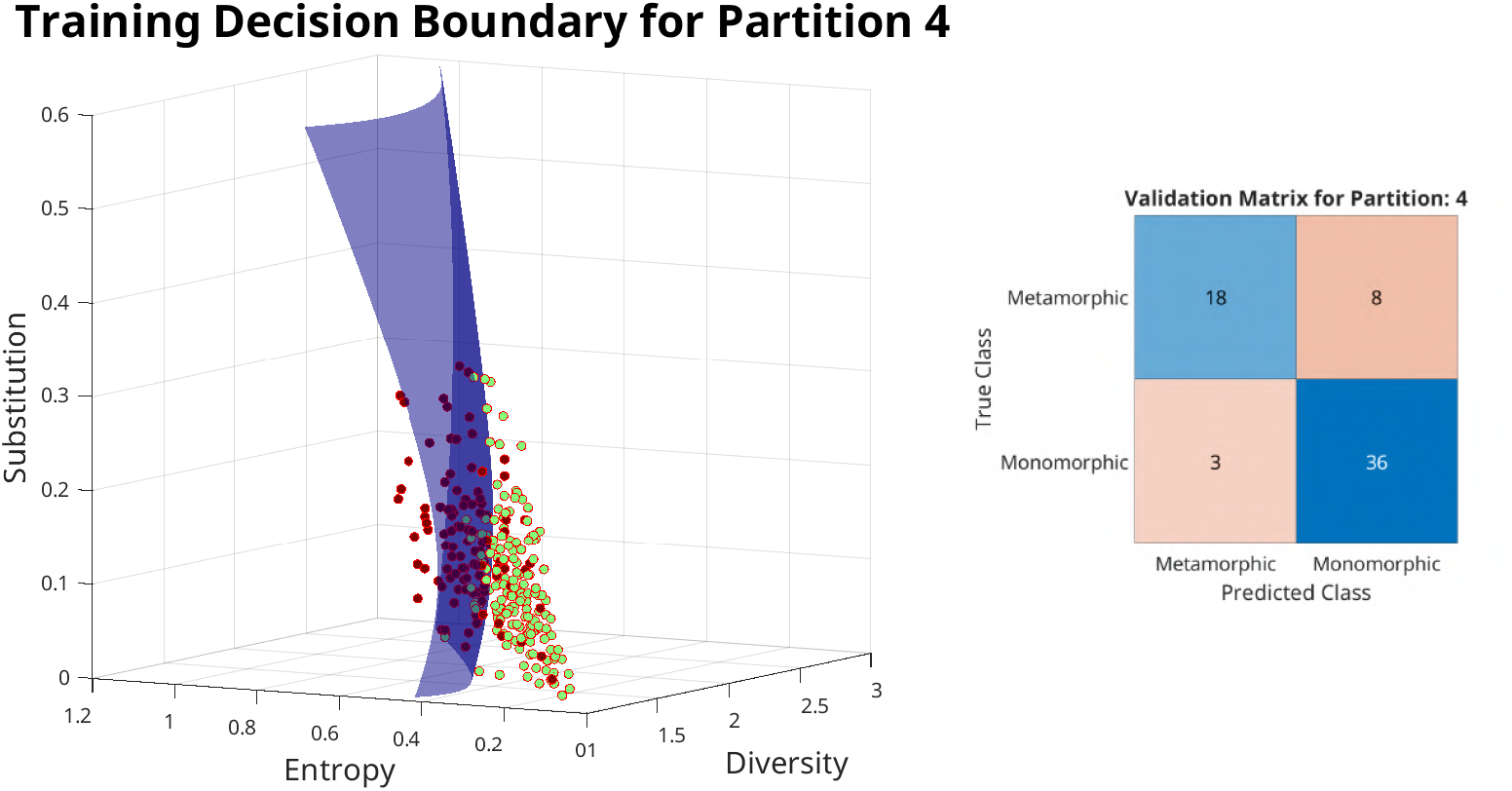
Decision boundary and the corresponding validation confusion matrix for the fourth training split partition

**Figure S3.e:**
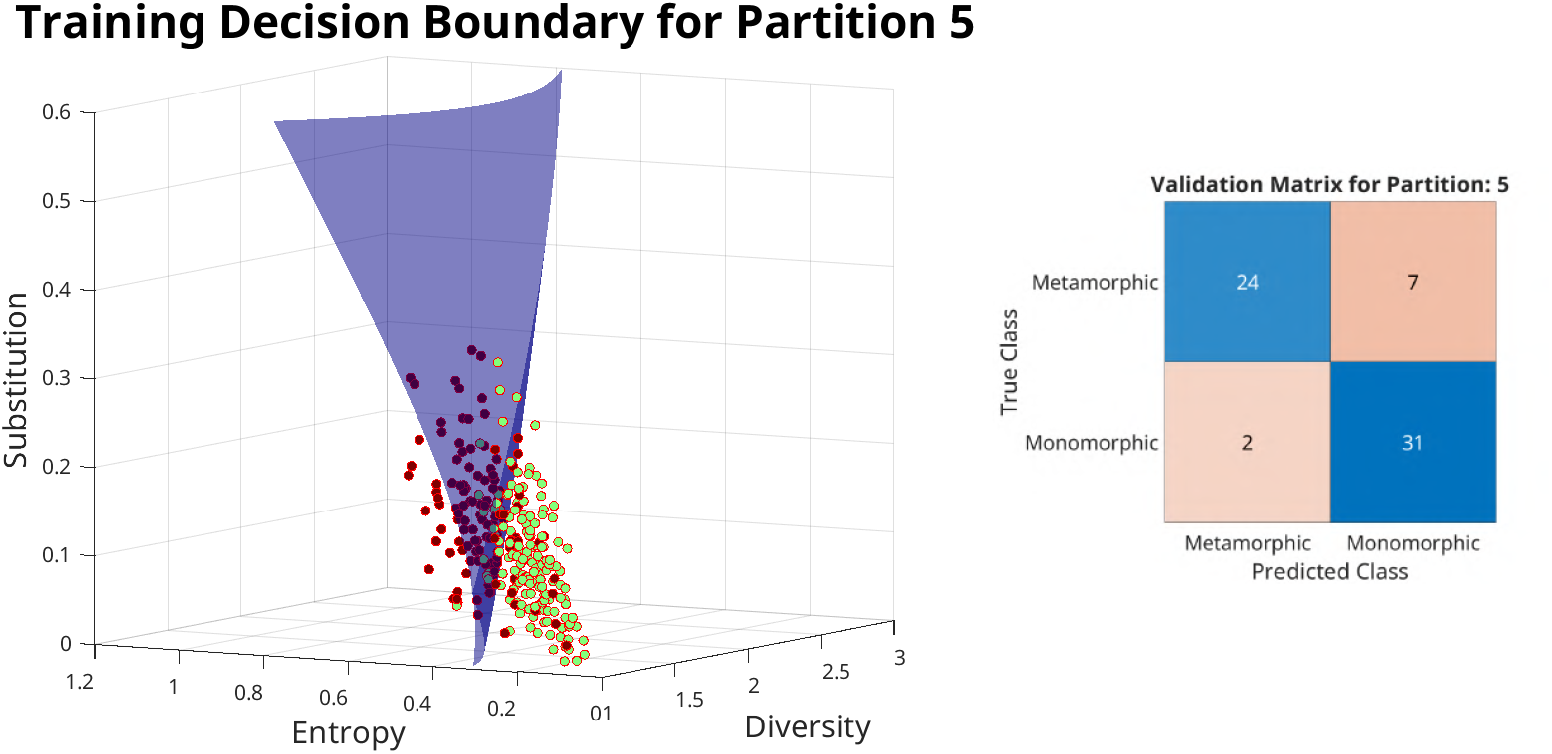
Decision boundary and the corresponding validation confusion matrix for the fifth training split partition

**Figure S3.f:**
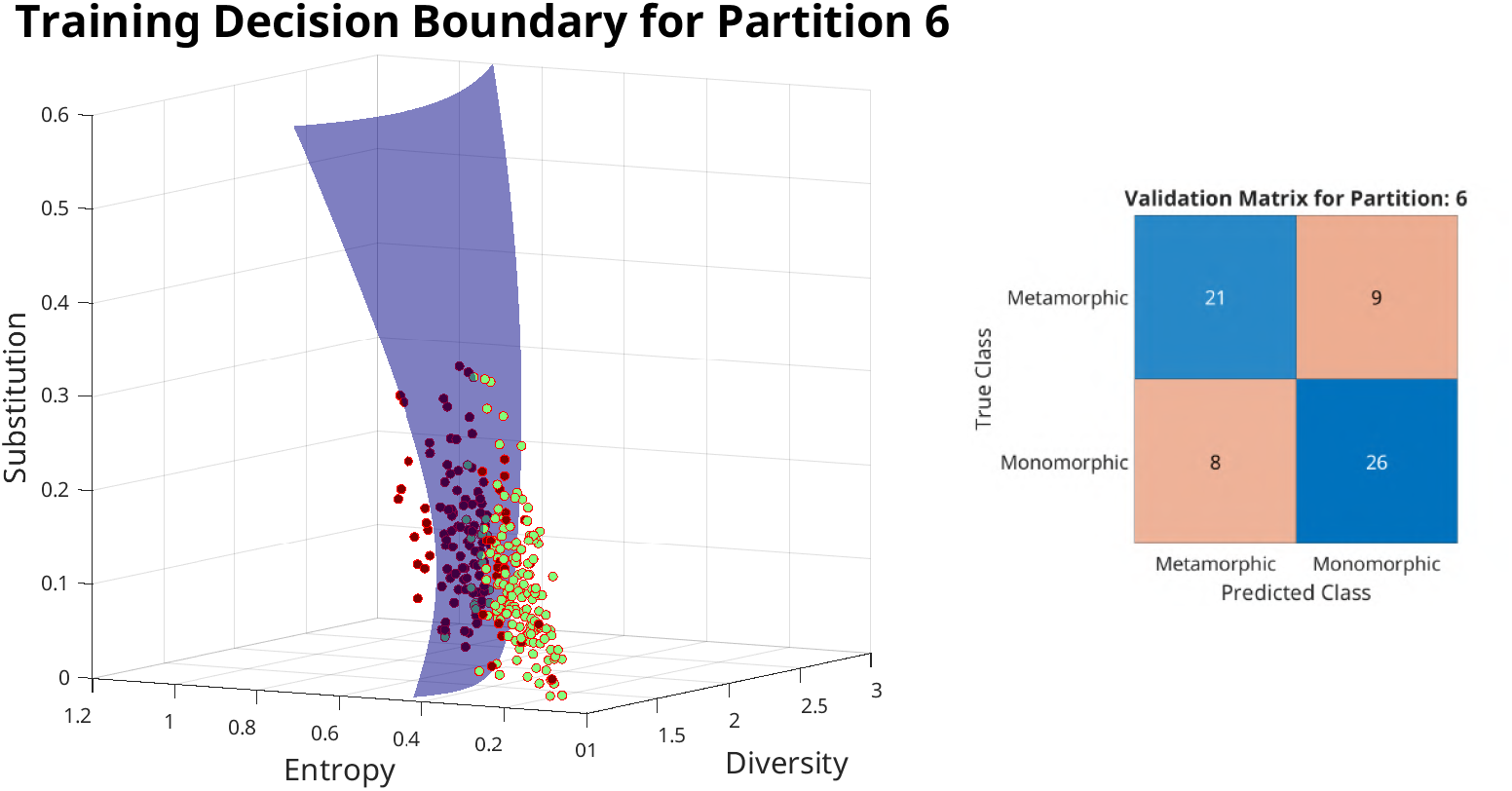
Decision boundary and the corresponding validation confusion matrix for the sixth training split partition

**Figure S3.g:**
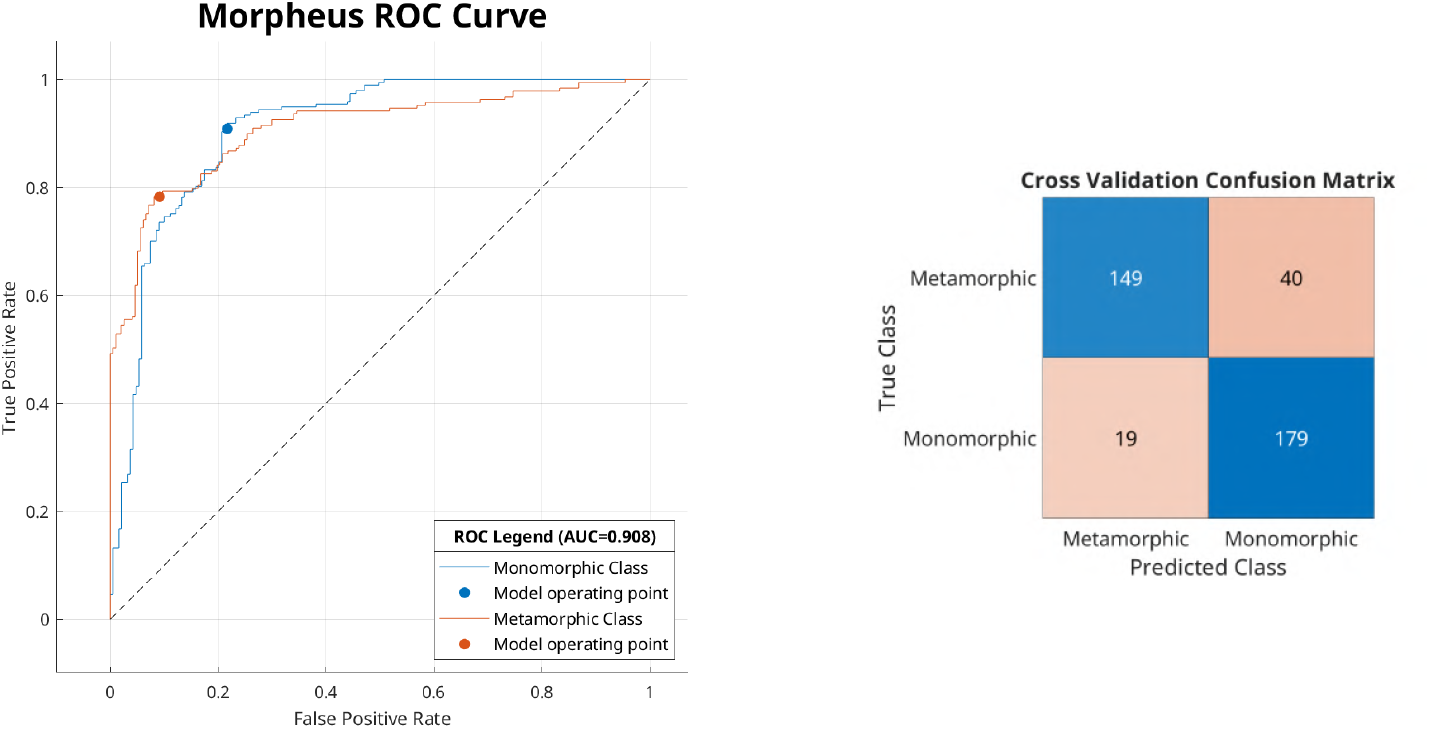
The Receiver-operating characteristic (ROC) curve for the final quadratic SVM classifier model and the final cross-validation matrix. The ROC curve is plotted considering the metamorphic protein class as the positive class. The area under the curve (AUC) for ROC curve is 0.89.

**Figure S3.h:**
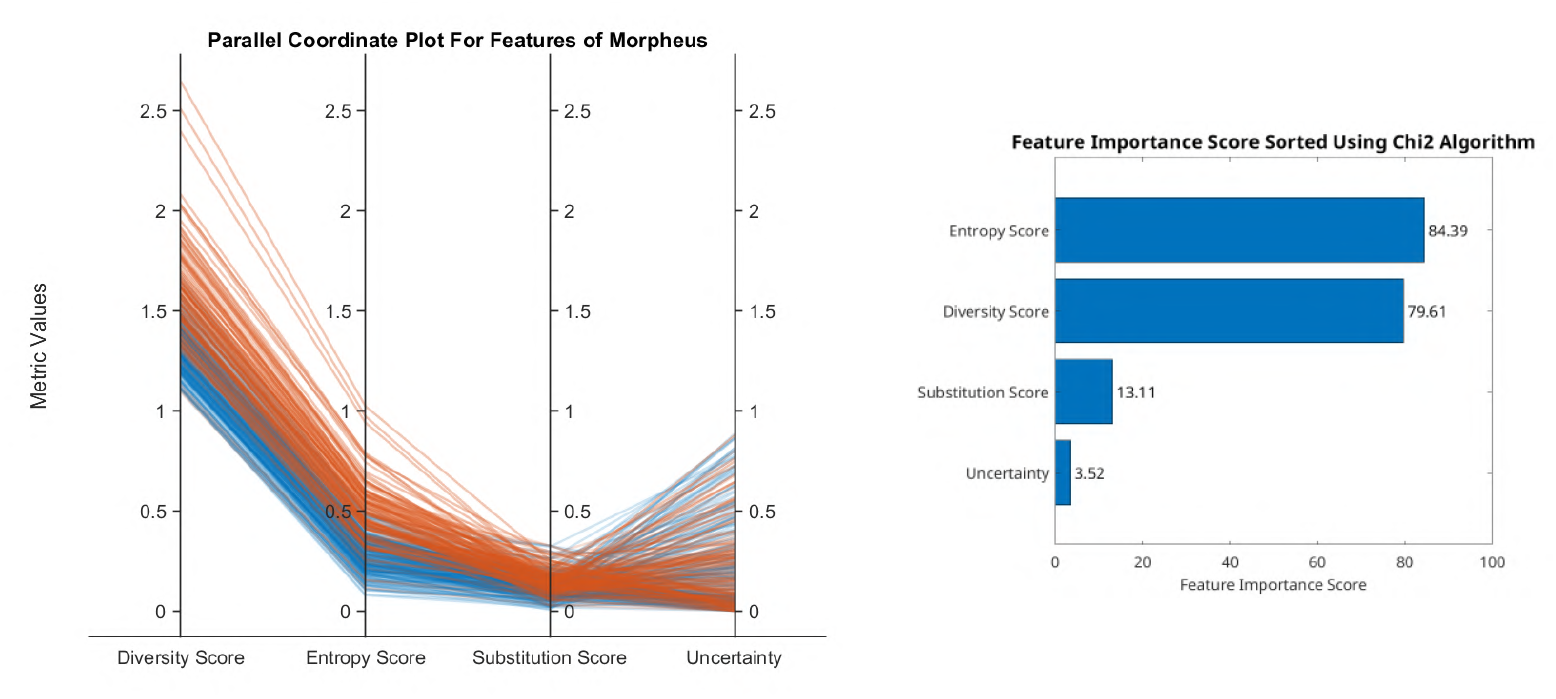
The parallel coordinate plot and *χ*^2^ scores for the four features diversity, entropy, substitution score and uncertainty. The lines plotted in orange represent metamorphic proteins, and the lines in blue represent monomorphic proteins in the training dataset. The features are not transformed while plotting but are shown as is.

#### 1.4 Figure S4: Proteome Data

Figures S4 shows the proteome data plotted on the feature space of diversity, entropy and substitution scores. The region to the left of the decision boundary represents the proteins that are predicted to be metamorphic, and the regions on the right are the ones predicted to be monomorphic.

**Figure S4.a:**
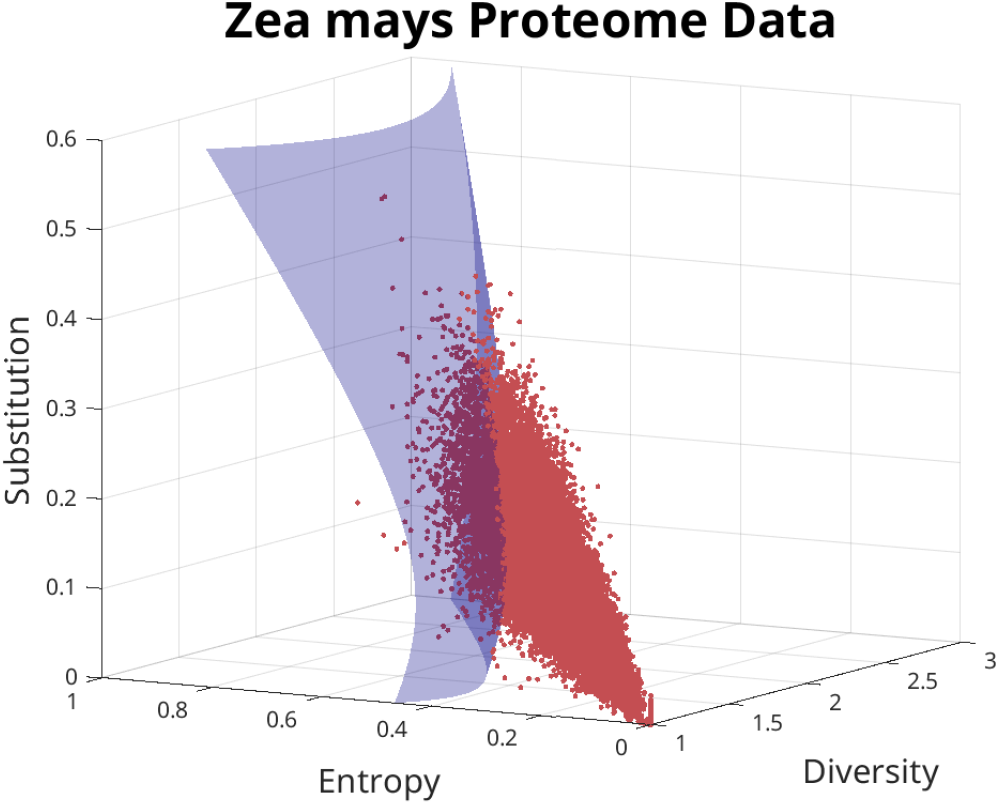
**Maize** proteome data plotted in the feature space of Diversity score, entropy score and substitution score. The surface represents the decision boundary obtained by the quadratic SVM model.

**Figure S4.b:**
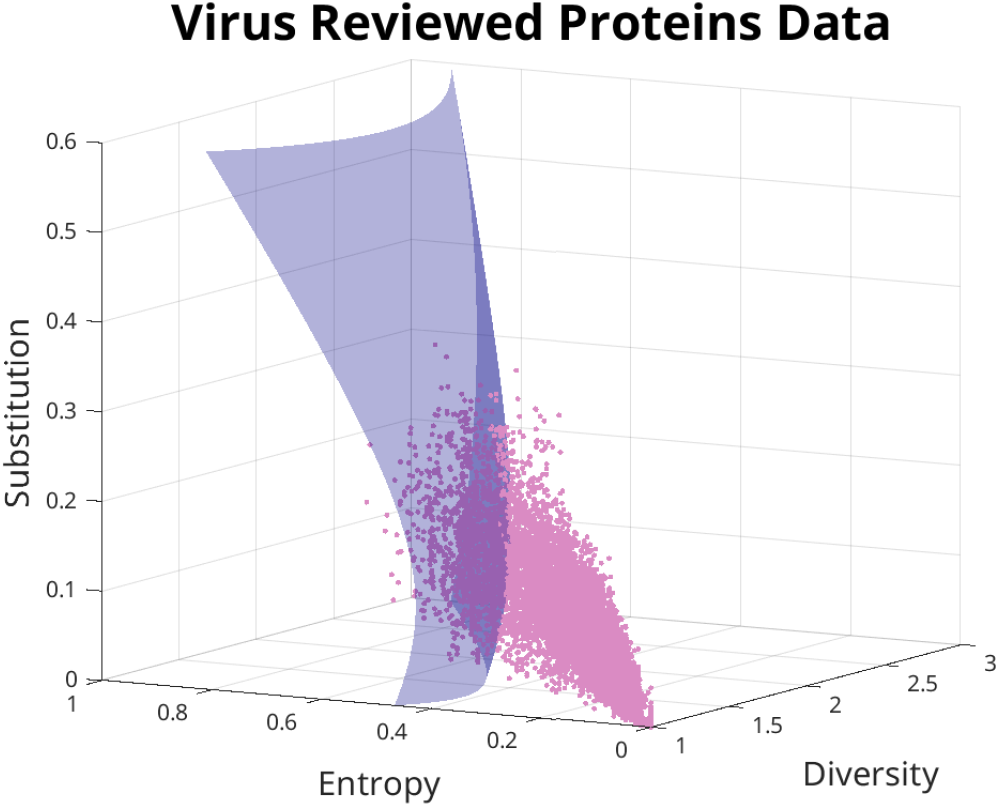
Manually reviewed proteins that belong to the viral proteomes are plotted in the feature space of Diversity score, entropy score and substitution score. The surface represents the decision boundary obtained by the quadratic SVM model.

#### 1.5 Figure S5: New Predictions

In the set of figures S5, the diversity metrics are plotted for the selected few predictions listed in the paper. The plots contain calculated helix and sheet propensities, the entropy scores, the diversity scores and Substitution scores at each sliding window. The uncertainty score at each fragment is also taken into a rolling average and overlayed on the diversity score.

**Figure S5.a:**
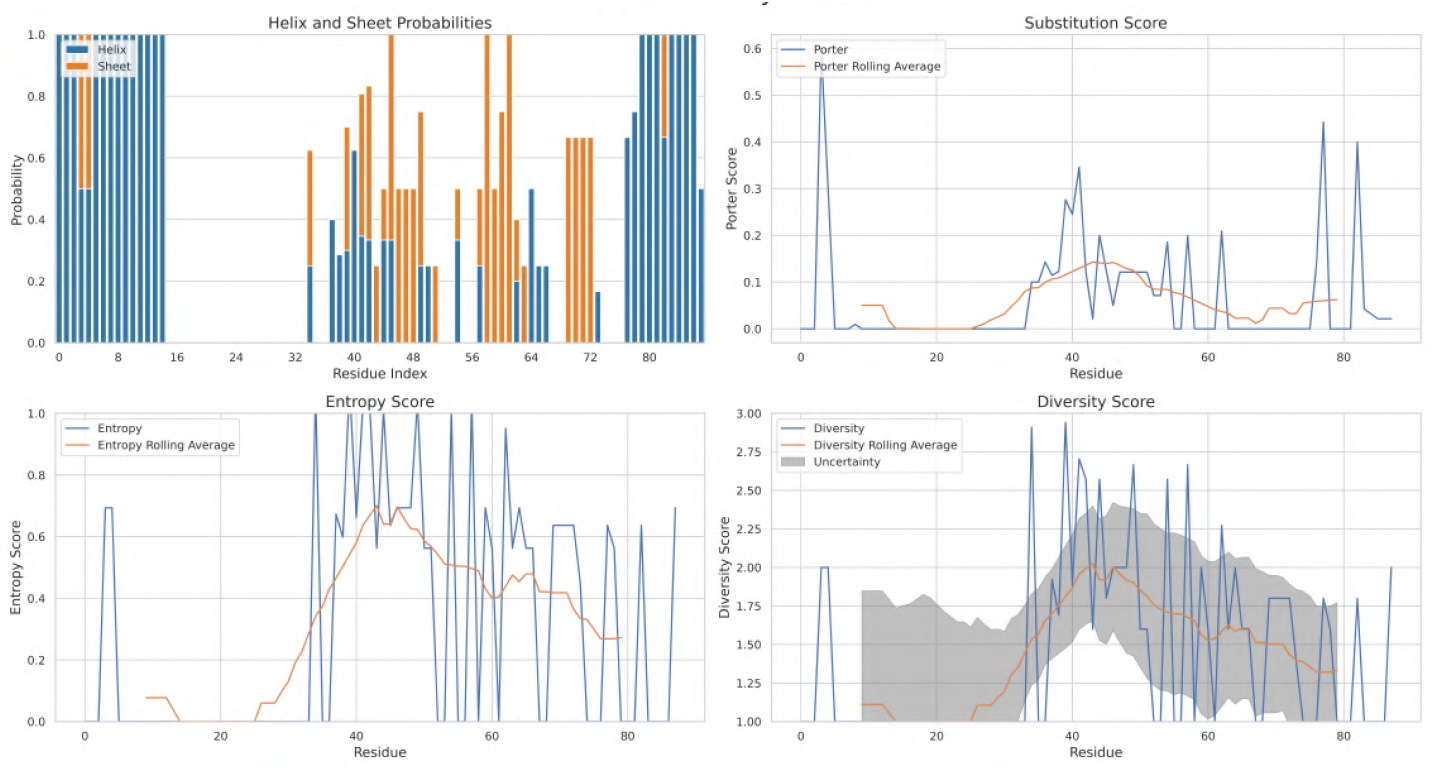
CXCL11: The diversity metrics plotted for the protein “CXCL11". Entropy, Diversity, Substitution and uncertainty are plotted above.

**Figure S5.b:**
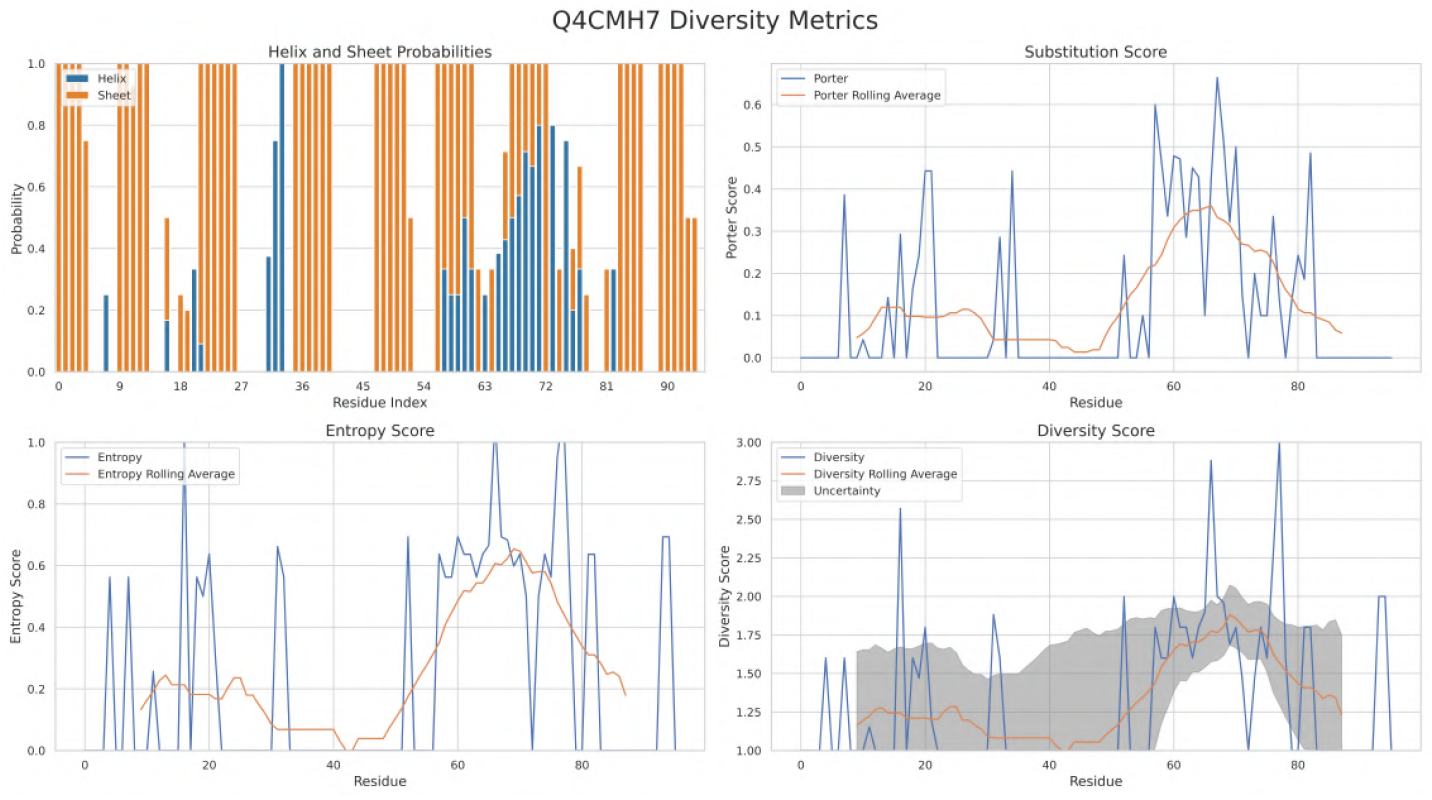
Guanine nucleotide-binding protein subunit beta-like protein: The diversity metrics plotted for the protein “Guanine nucleotide-binding protein subunit beta-like protein". Entropy, Diversity, Substitution and uncertainty are plotted above.

**Figure S5.c:**
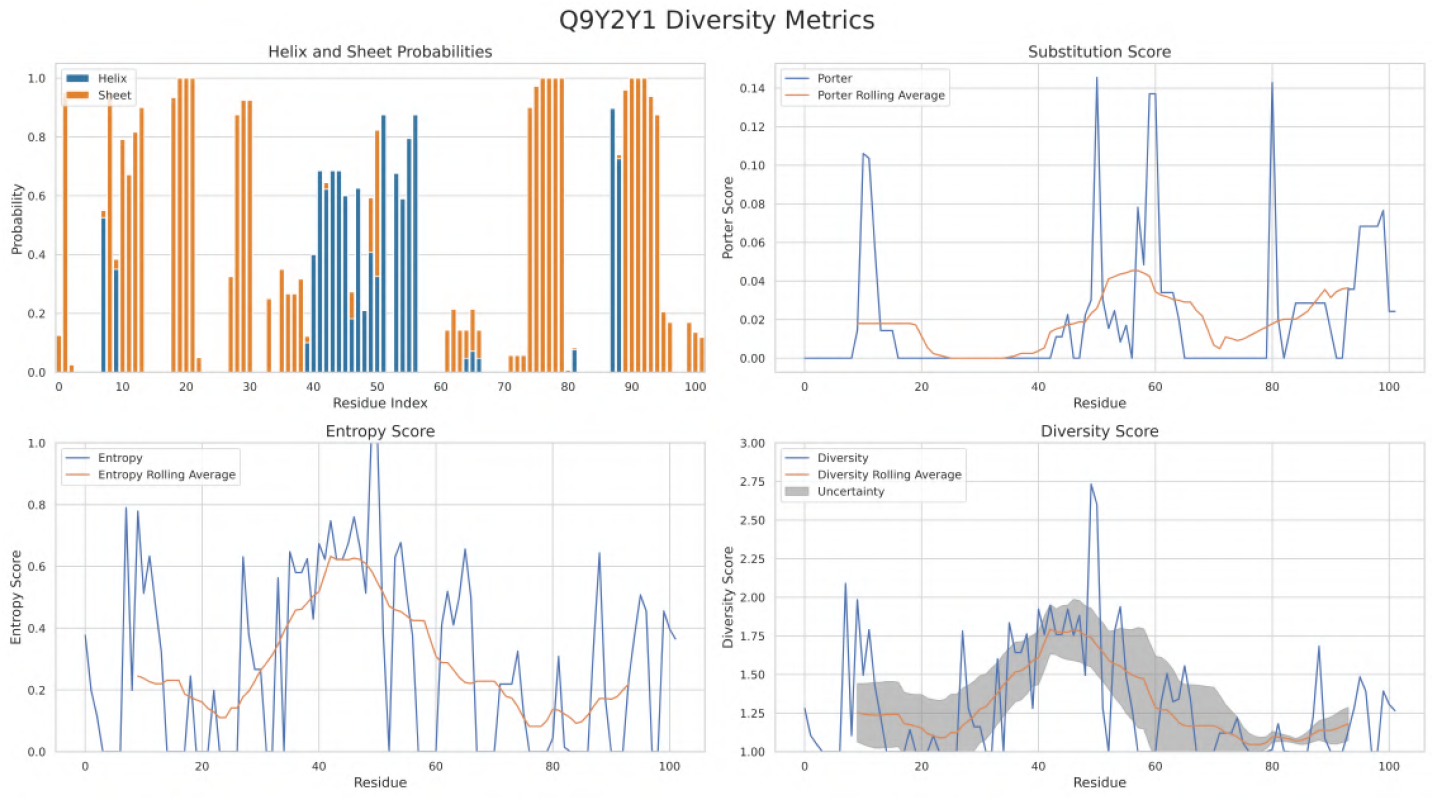
DNA-directed RNA polymerase III subunit RPC10: The diversity metrics plotted for the protein “DNA-directed RNA polymerase III subunit RPC10". Entropy, Diversity, Substitution and uncertainty are plotted above.

**Figure S5.d:**
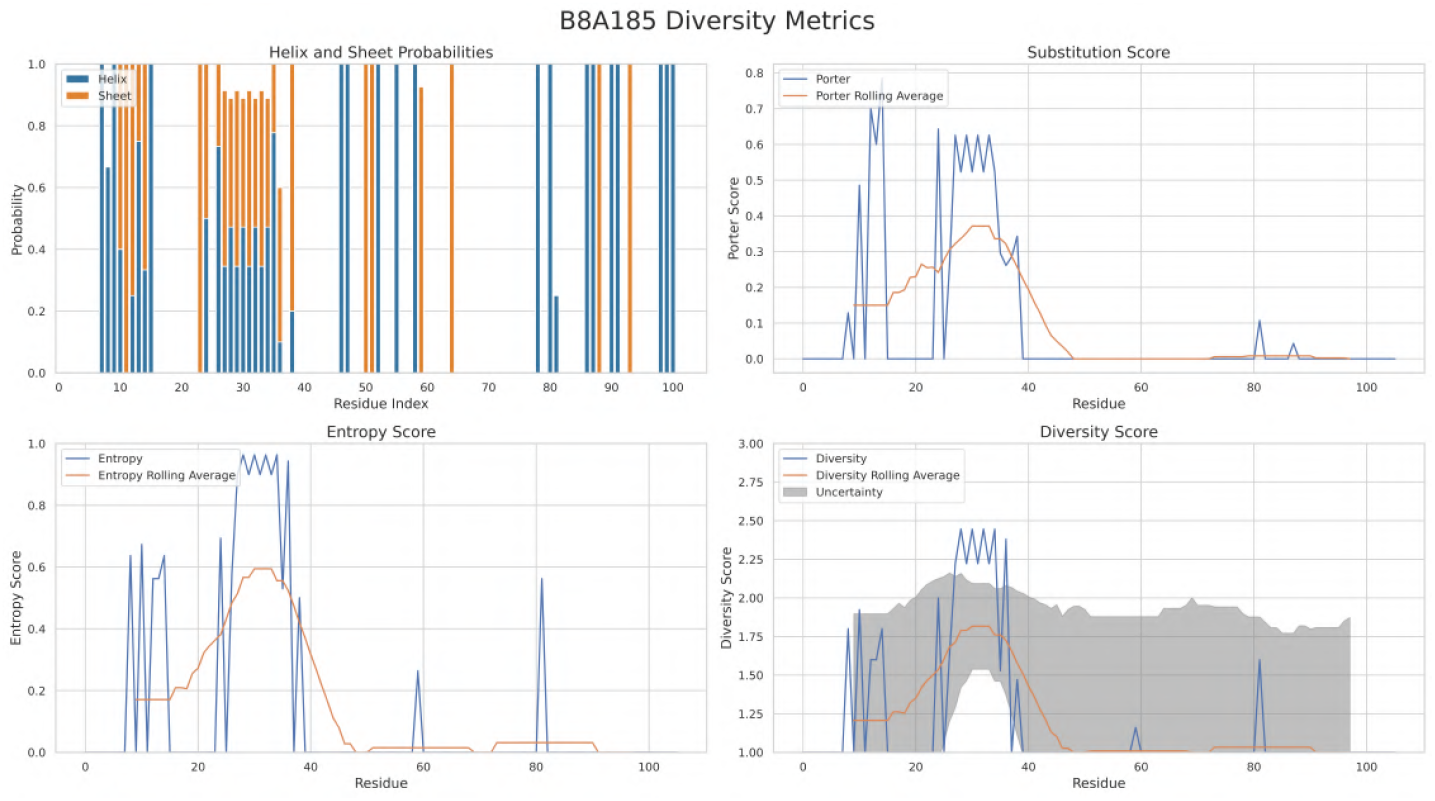
Secreted protein: The diversity metrics plotted for the Secreted protein. Entropy, Diversity, Substitution and uncertainty are plotted above.

**Figure S5.e:**
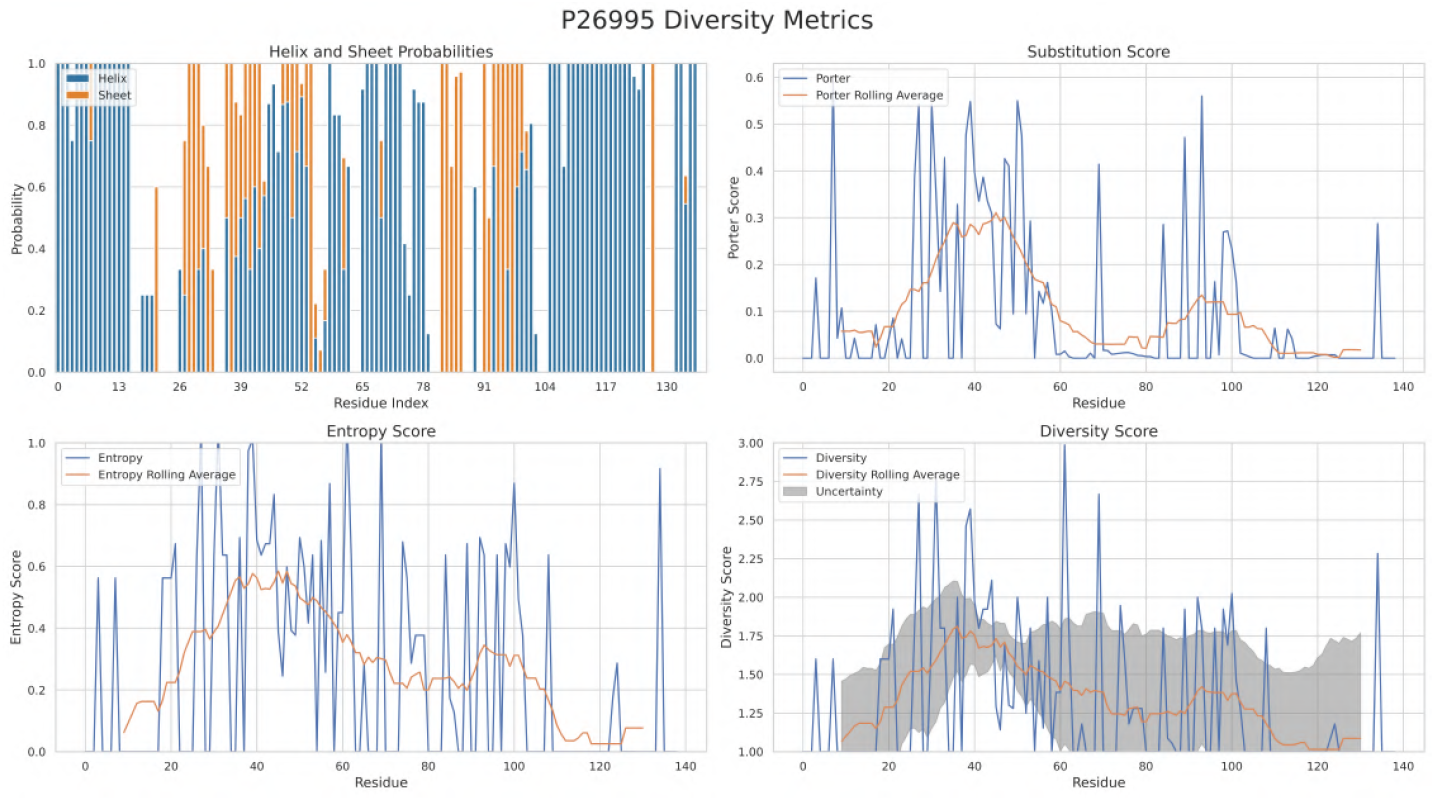
Transcriptional anti-antiactivator ExsC: The diversity metrics plotted for the protein “Transcriptional anti-antiactivator ExsC". Entropy, Diversity, Substitution and uncertainty are plotted above.

**Figure S5.f:**
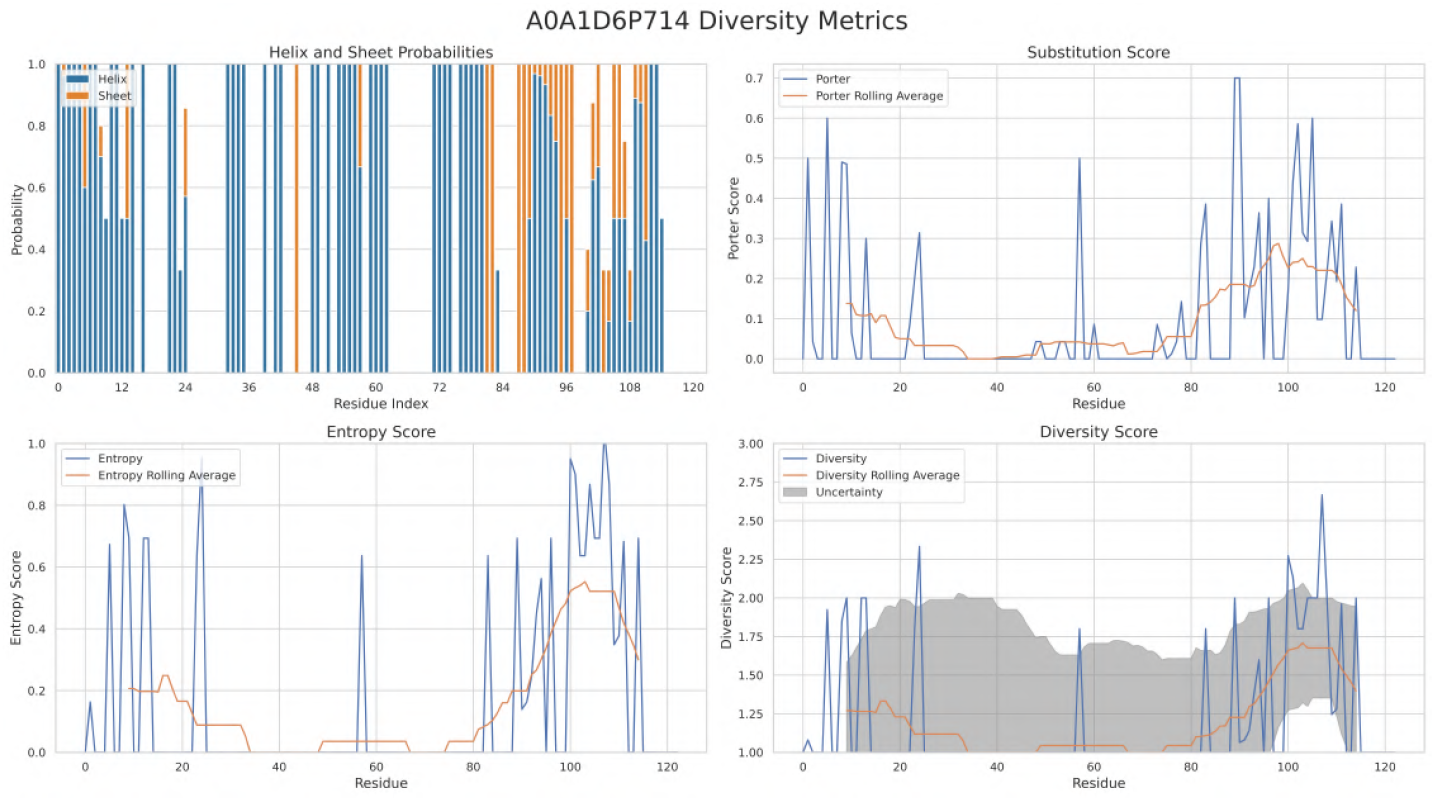
Extensin-like protein: The diversity metrics plotted for the protein “Extensin-like protein". Entropy, Diversity, Substitution and uncertainty are plotted above.

**Figure S5.g:**
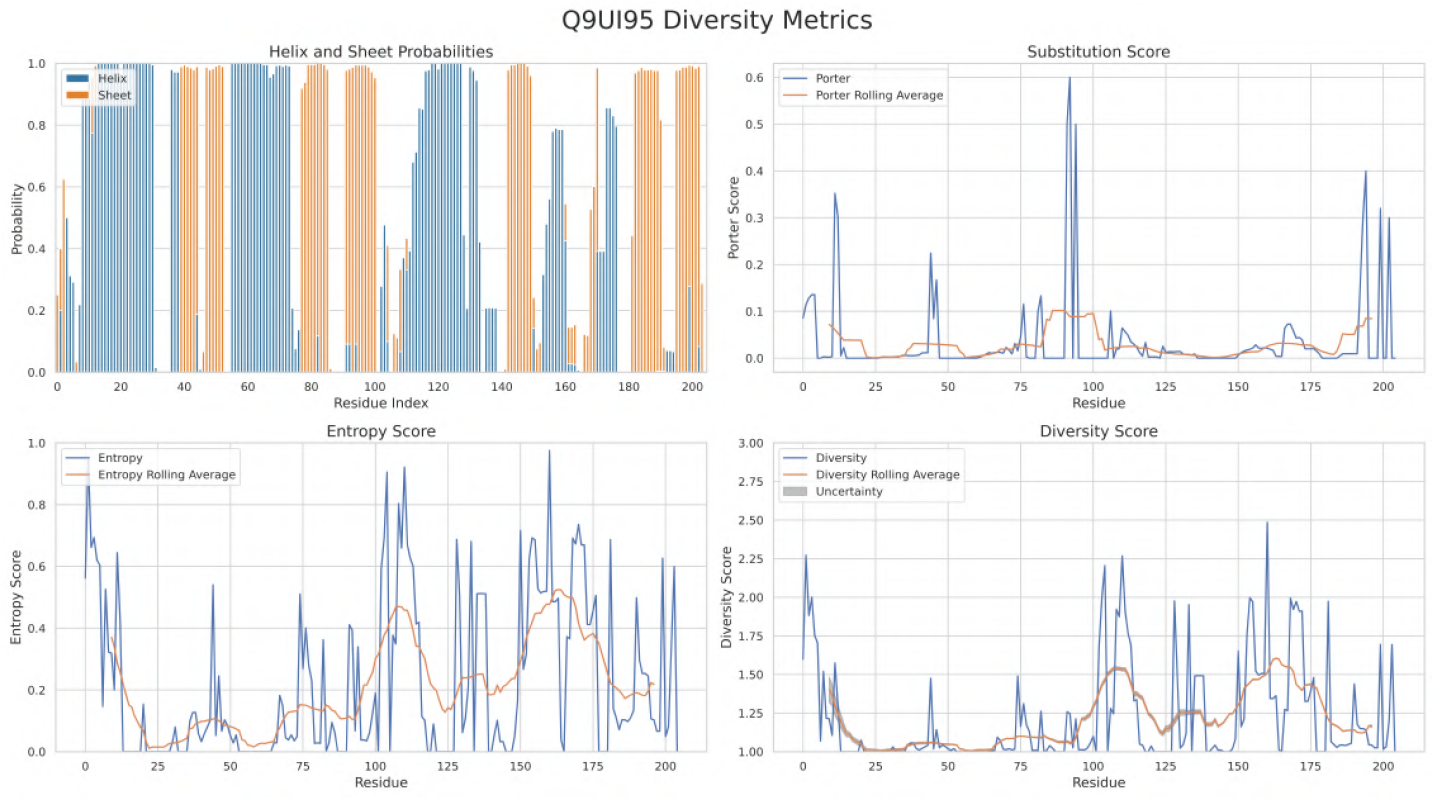
Mitotic spindle assembly checkpoint protein MAD2B: The diversity metrics plotted for the protein “Mitotic spindle assembly checkpoint protein MAD2B". Entropy, Diversity, Substitution and uncertainty are plotted above.

**Figure S5.h:**
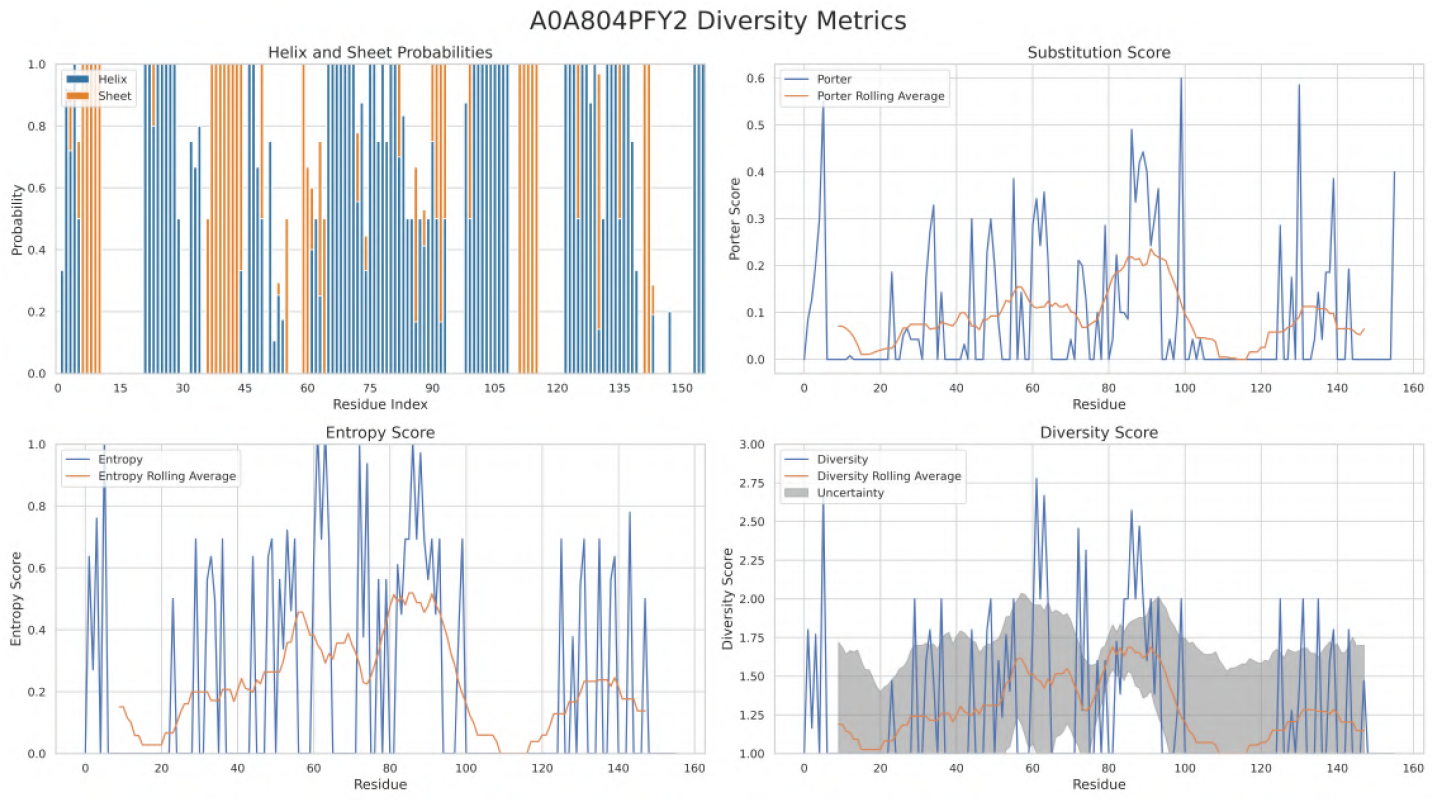
Photolyase/cryptochrome alpha/beta domain-containing protein: The diversity metrics plotted for the protein “Photolyase/cryptochrome alpha/beta domain-containing protein". Entropy, Diversity, Substitution and uncertainty are plotted above.

**Figure S5.i:**
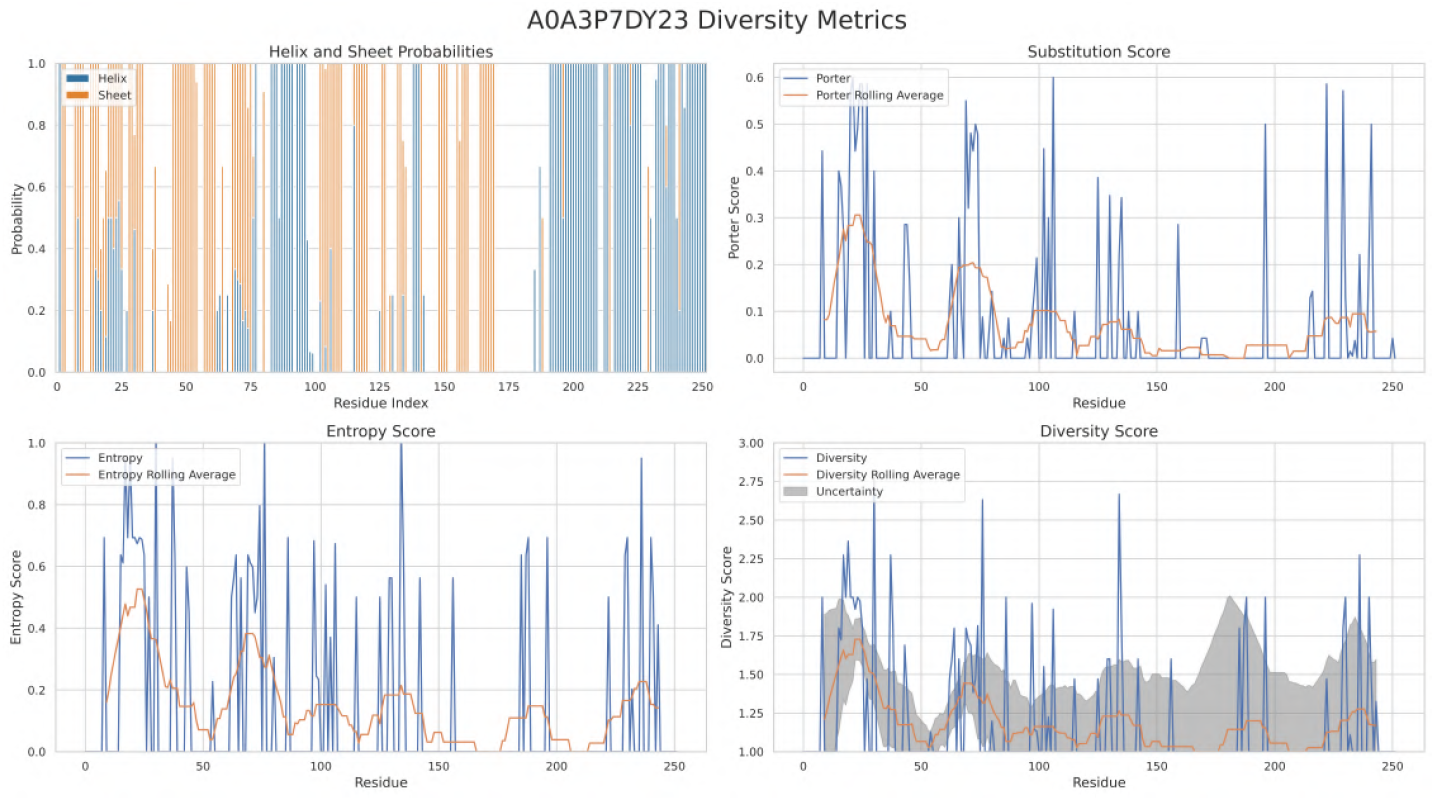
S1 motif domain-containing protein: The diversity metrics plotted for the protein “S1 motif domain-containing protein". Entropy, Diversity, Substitution and uncertainty are plotted above.

**Figure S5.j:**
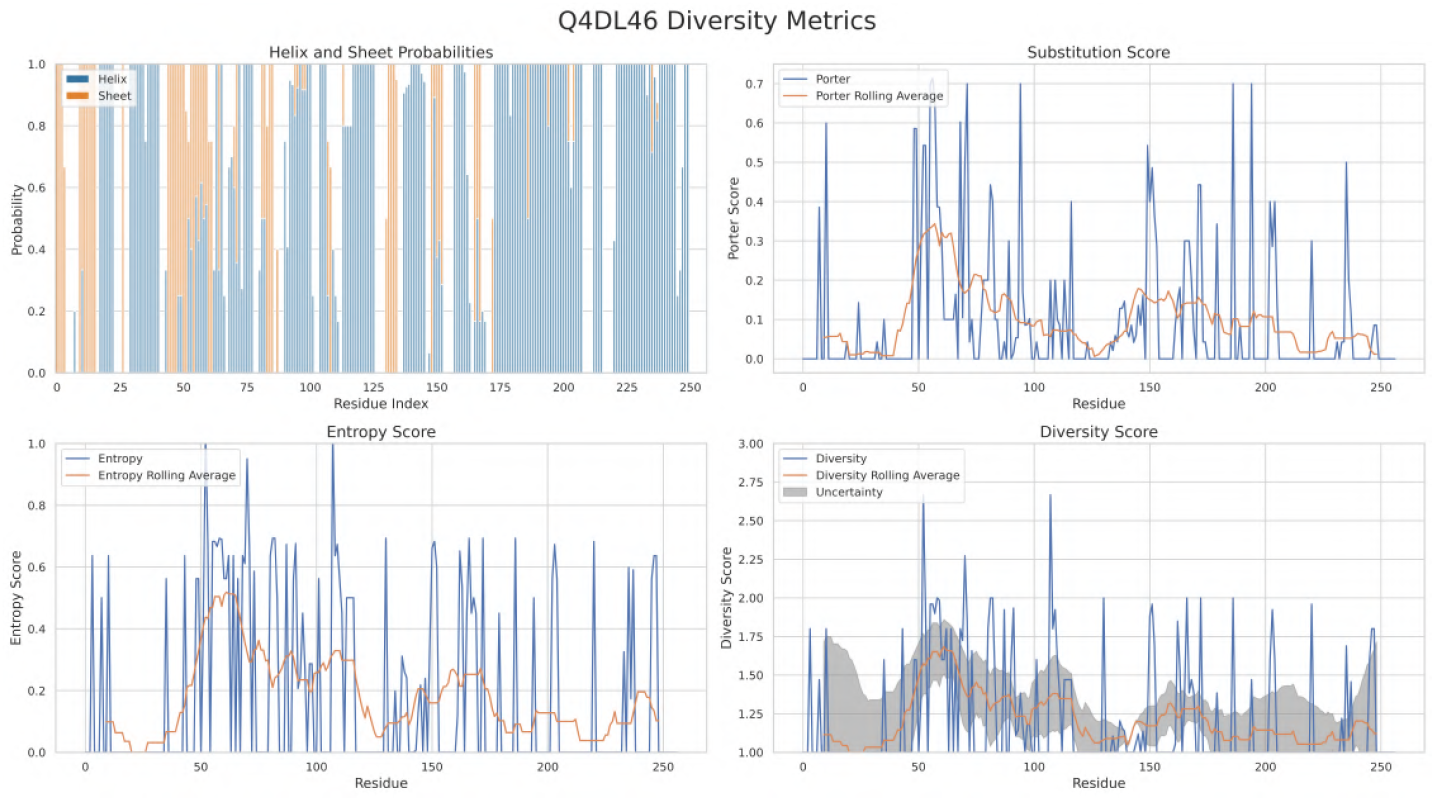
Copper-transporting ATPase-like protein, putative: The diversity metrics plotted for the protein “Copper-transporting ATPase-like protein, putative". Entropy, Diversity, Substitution and uncertainty are plotted above.

### 2 Dataset

The data used for training the SVM model is borrowed from existing literature on fold-switching proteins and is provided in the tables below. In table S1, the columns PDB1 and PDB2 represent the PDB ID along with the chain ID for the two distinct secondary structures that were solved for the fold-switching protein.

#### 2.1 Fold-switching Proteins Dataset

**Table S1:**
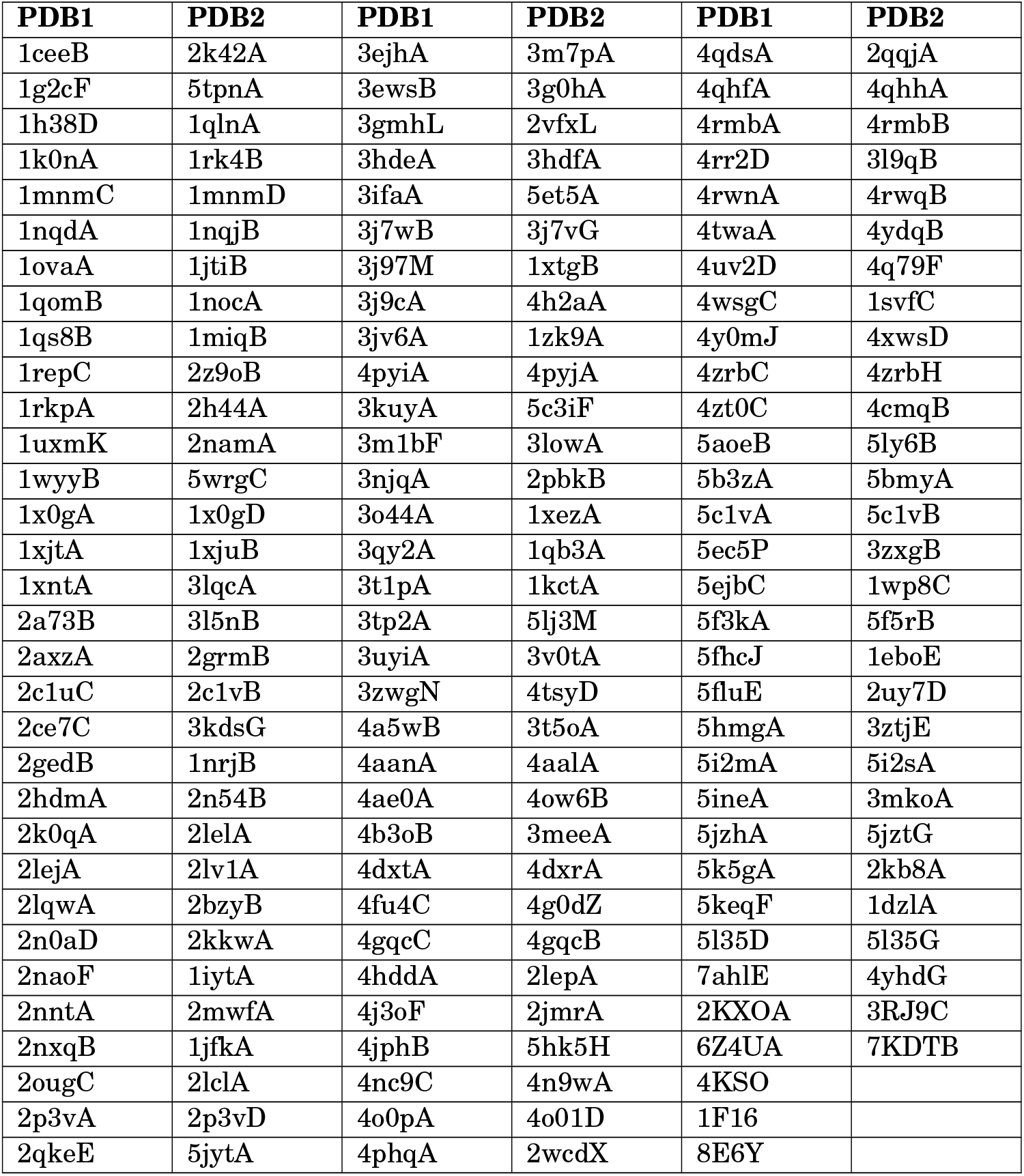
189 Metamorphic proteins that are used for training the model. PDB1 and PDB2 refer to the two different solved structures of the protein

#### 2.2 Monomorphic Proteins Dataset

**Table S2:**
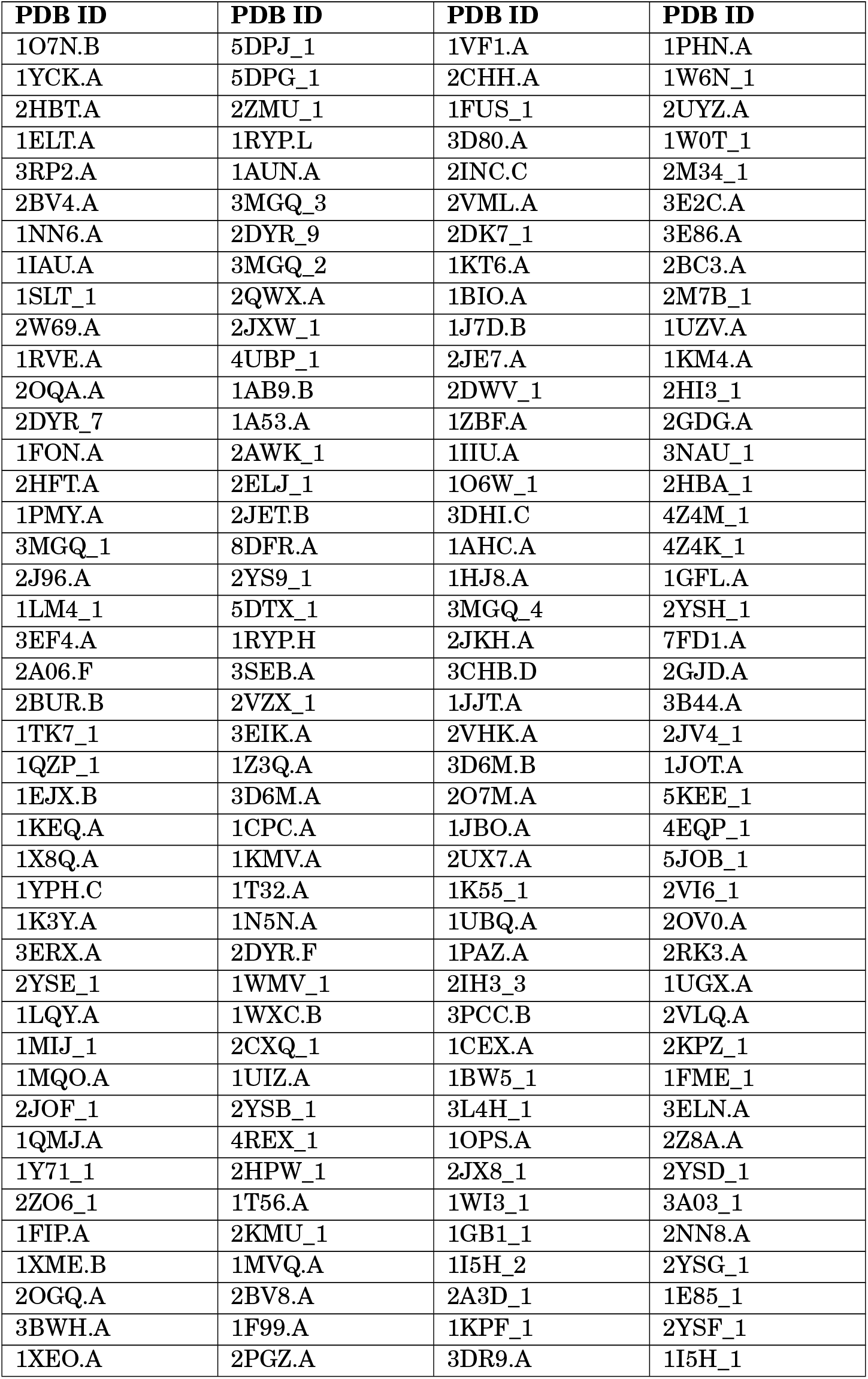
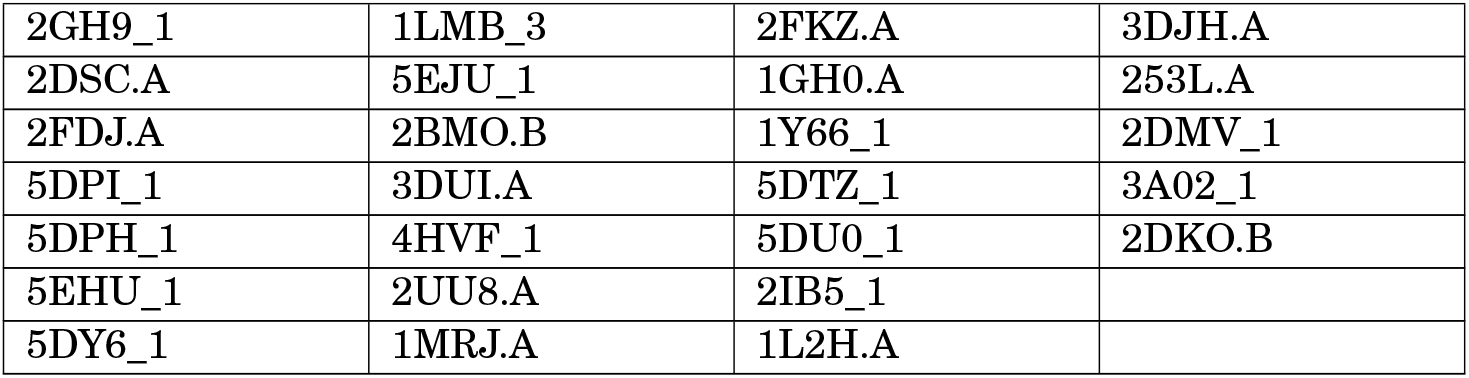
198 Monomorphic proteins that are used for training the model. The character after the dot/underscore refers to the chain ID of the protein

### 3 Optimizations

#### a. Rolling window width

The Rolling average window width is optimized by considering a range of values from 5 to 25 and picking the one that gives the best Mattew’s Correlation coefficient (MCC) when applied to the training dataset. A six-fold cross-validation scheme is applied when the MCC value is calculated with each of the window widths. The validation accuracy and MCC score are calculated, and the window width that gives the best MCC score is selected. Here the columns are window width, Accuracy when a quadratic kernel is used, MCC when a quadratic kernel is used, Accuracy when a linear kernel is used, MCC when a linear kernel is used for the SVM.

**Figure S5.k:**
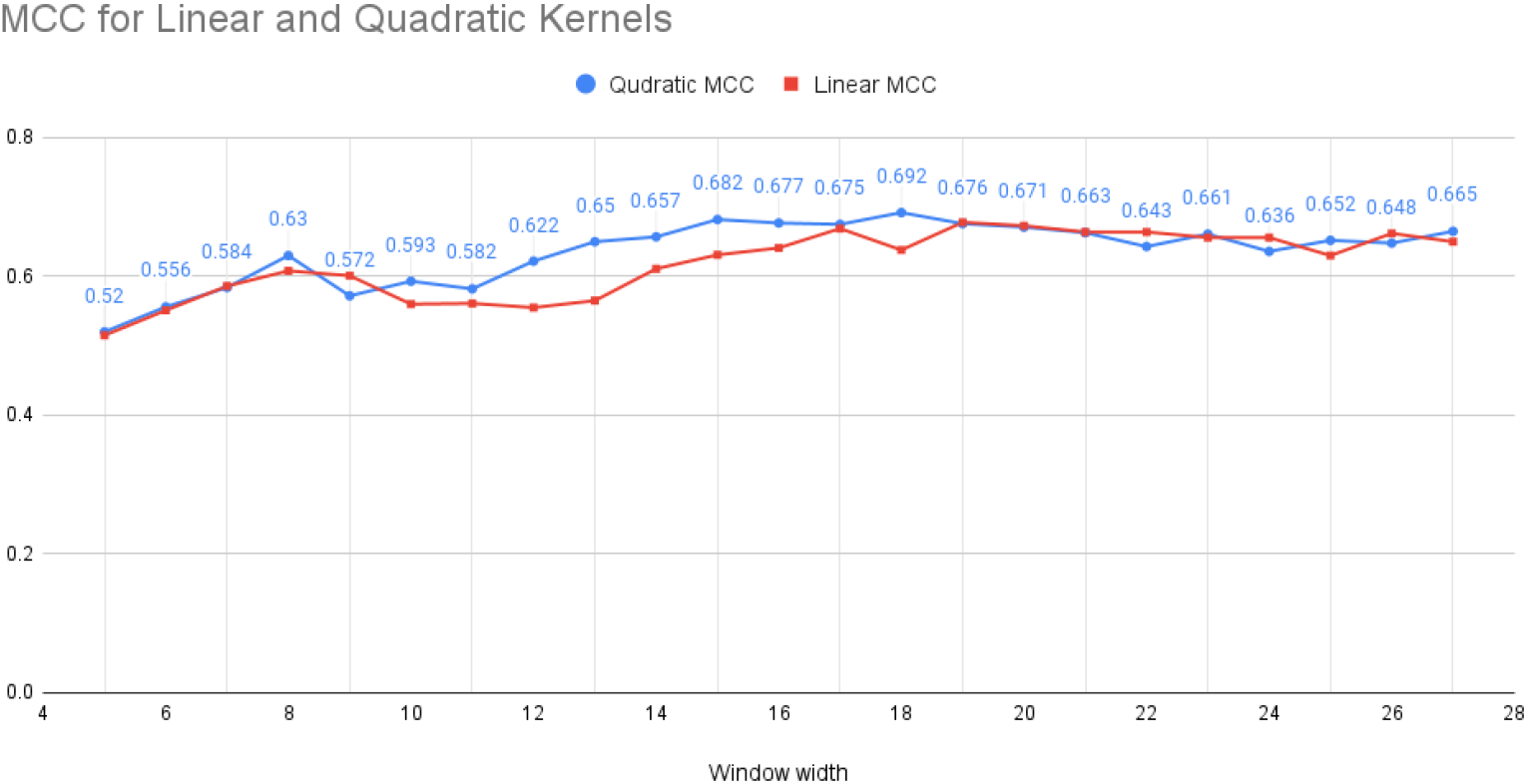
The MCC values for each window width is plotted and the top two window widths (8,18) are selected later to be used for calculating the diversity metrics.

#### b. Substitution Matrix

Similarly to optimizing the window width, the substitution matrix is optimized using a range of parameters the form shown below:

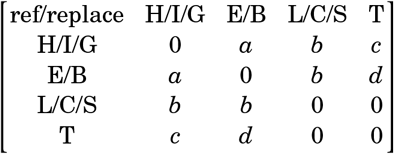

**Table S3:** The Accuracy and MCC measures are provided for each of the value of averaging window width ranging from 5 to 27. These measures are calculated by performing a 6-fold cross-validation on the training dataset.

| Window width | Quad Acc | Quad SVM MCC | Lin SVM Acc | Lin SVM MCC |
| --- | --- | --- | --- | --- |
| 5 | 77.2 | 0.52 | 75.1 | 0.515 |
| 6 | 77.8 | 0.556 | 77.5 | 0.551 |
| 7 | 79.1 | 0.584 | 79.3 | 0.586 |
| 8 | 81.4 | 0.63 | 81.5 | 0.608 |
| 9 | 78.5 | 0.572 | 80.1 | 0.601 |
| 10 | 79.6 | 0.593 | 78 | 0.56 |
| 11 | 78.8 | 0.582 | 78 | 0.561 |
| 12 | 80.8 | 0.622 | 77.7 | 0.555 |
| 13 | 82.1 | 0.65 | 78.2 | 0.565 |
| 14 | 82.6 | 0.657 | 80.57 | 0.611 |
| 15 | 83.9 | 0.682 | 81.6 | 0.631 |
| 16 | 83.6 | 0.677 | 82.1 | 0.641 |
| 17 | 83.6 | 0.675 | 83.4 | 0.669 |
| 18 | 84.4 | 0.692 | 81.8 | 0.638 |
| 19 | 83.6 | 0.676 | 83.9 | 0.678 |
| 20 | 83.4 | 0.671 | 83.6 | 0.673 |
| 21 | 82.9 | 0.663 | 83.1 | 0.664 |
| 22 | 81.8 | 0.643 | 83.1 | 0.664 |
| 23 | 82.8 | 0.661 | 82.8 | 0.656 |
| 24 | 81.5 | 0.636 | 82.8 | 0.656 |
| 25 | 82.3 | 0.652 | 81.5 | 0.63 |
| 26 | 82.3 | 0.648 | 83.1 | 0.662 |
| 27 | 83.1 | 0.665 | 82.6 | 0.65 |

The values for a,b,c and d were sampled first in a coarse manner, where the values of all the 4 parameters range from 0 to 1 with a step of 0.2 for each one. Then once a substitution matrix that gave the maximum MCC value for the substitution score is found, the sampling is made much finer with a step value of 0.05 for each of the parameter. That way we arrived at a final substitution matrix as shown (d=0):

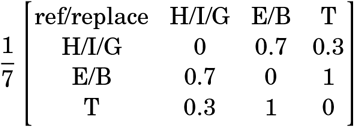

#### c. Fragment Size

The fragment size is optimized via trial and error. For the training dataset. Fragment picking was performed on the training dataset with the fragment sizes: 6, 7 and 8. When fragment picking is performed with a fragment size of 8, the number of hits, i.e. proteins containing the identical fragment, was quite low with the database currently in use. While this does not give us enough data to work with, using fragment size 6 incurred the problem of the fragment secondary structure diversity, being all over the place. The 6-mers did not seem to truly represent the diversity taken by the protein in question. Therefore the fragment size is set to 7 and the high accuracy in prediction seems to validate the choice of picking 7-mers.

#### d. Middle Residue

After fragment picking, the residue that was taken into consideration for the secondary structure analysis is the middle residue (4th residue of the 7-mer fragments). But upon considering other residues, there was a small change in accuracy measures. Although the accuracy measures when whichever residue is taken into consideration did not vary beyond a significant number standard deviation, we still chose to work with the one that gave the highest MCC score. Table S4 shows the variation of accuracy when the fragment in consideration is changed from the 2nd residue to the 6th residue of the 7-mer. Accordingly, we chose the final model to work with the 3rd residue.

**Table S4:** The Accuracy measures for the residue in consideration starting from the 2nd residue to the 6th residue of the 7-mer fragment.

| Measure | 2nd RSD | 3rd RSD | 4th RSD | 5th RSD | 6th RSD |
| --- | --- | --- | --- | --- | --- |
| TPR | 0.82 (0.08) | 0.788 (0.08) | 0.777 (0.05) | 0.874 (0.04) | 0.787 (0.02) |
| SPC | 0.852 (0.06) | 0.904 (0.06) | 0.863 (0.07) | 0.737 (0.12) | 0.886 (0.05) |
| PRE | 0.844 (0.05) | 0.887 (0.08) | 0.845 (0.07) | 0.771 (0.08) | 0.872 (0.05) |
| ACC | 83.42 (4.3%) | 84.75 (6.0%) | 81.87 (4.4%) | 80.55 (6.3%) | 83.93 (2.7%) |
| F1 | 0.829 (0.04) | 0.834 (0.08) | 0.808 (0.04) | 0.818 (0.05) | 0.827 (0.02) |
| MCC | 0.6727 (0.08) | 0.6983 (0.12) | 0.6389 (0.09) | 0.6185 (0.12) | 0.678 (0.05) |

#### e. Support Machine Vector Model

An SVM model with a quadratic kernel is chosen for making predictions on data the training model has not encountered before. Support vector machine is chosen as they achieve good performance on many classification tasks. Kernels make SVMs more flexible and able to handle nonlinear problems, and here as we are using a quadratic kernel, we do not have to worry about the model complexity. As we have a relatively small dataset to train with, this restricts the choices of many machine learning models we could use. A quadratic SVM model gives us good accuracy while still maintaining a low model complexity.

The hyperparameters box constraint and Kernel scale were optimized using a Bayesian optimization technique in MATLAB.

